# Gating current noise produced by Brownian models of a voltage sensor

**DOI:** 10.1101/2021.01.13.426543

**Authors:** Luigi Catacuzzeno, Fabio Franciolini, Francisco Bezanilla, Robert S. Eisenberg

**Author notes:** **Correspondence to:** Luigi Catacuzzeno Department of Chemistry, Biology and Biotechnology University of Perugia, via Elce di Sotto 8, 06123 Perugia - Italy, **Email address**.

## Abstract

The activation of voltage-dependent ion channels is associated with the movement gating charges, that give rise to gating currents. Although gating currents originating from a single channel are too small to be detected, analysis of the fluctuations of macroscopic gating currents originating from a population of channels can make a good guess of their magnitude. The analysis of experimental gating current fluctuations, when interpreted in terms of a Markov model of channel activation, are in accordance with the presence of a main step along the activation pathway carrying 2.3-2.4 e_0_ of charge. To give a physical interpretation to these results and to relate them to the known atomic structure of the voltage sensor domain, we employed a Brownian model of voltage-dependent gating that we recently developed using structural information and applying the laws of electrodynamics. The model was capable to reproduce gating currents and gating current fluctuations essentially similar to those experimentally observed. The detailed study of this model output, also performed by making several simplifications aimed at understanding the basic dependencies of the gating current fluctuations, suggests that in real ion channels the voltage sensor does not move in a fully Markovian regimen due to the relatively low (<5 kT) energy barriers separating successive intermediate states. As a consequence, the simultaneous jump of multiple gating charges through the gating pore becomes frequent, and this occurrence is at the origin of the relatively high single-step charge detected by assuming Markovian behavior.

## Introduction

The activation of voltage-dependent Na_v_ and K_v_ channels in response to a depolarization is a vital step in the action potential, the long range signal of the nervous system and skeletal and cardiac muscle. This process produces a small but detectable component of current – the gating current – originating from the movement of protein charges, shown to be mostly associated with the voltage sensors for channels gating (Bezanilla, 2008, 2018). Experimental techniques today amplify the gating current by increasing the cell membrane density of voltage sensors enormously, and recording the macroscopic gating current.

Although transitions of individual voltage sensors do not generate experimentally detectable currents, they do produce noticeable fluctuations in the macroscopic gating current (Conti and Stühmer, 1989; Sigg, Stefani and Bezanilla, 1994). The analysis of the variance of these fluctuations allows estimates of the magnitude and time distribution of the elementary charge movements, when reasonable assumptions about the elementary charge movement are made, as done by (Conti and Stühmer, 1989) in Na_v_ and (Sigg, Stefani and Bezanilla, 1994) in Shaker K_v_. Both studies found that these fluctuations, assumed to originate from the individual movements of voltage sensors, produce a variance in measured current proportional to the mean current, a typical feature of shot noise (Crouzy and Sigworth, 1993; Landauer, 1993, 1996; Iannaccone *et al.*, 1998; Sigg, Qian and Bezanilla, 1999).

Shot noise is expected from traditional discrete state (Markov) models when the voltage sensor moves instantaneously from one stable position to the next, along the activation pathway (Crouzy and Sigworth, 1993). Notably, the interpretation of the experimental data on gating current fluctuation with Markov models suggests the presence of a main step along the activation pathway carrying a charge of 2.3-2.4e_0_ (Conti and Stühmer, 1989; Sigg, Stefani and Bezanilla, 1994).

Later studies performed with a drift-diffusion model of the voltage sensor showed that a charge diffusing along an energy profile produces a shot noise only if the energy profile encountered by the voltage sensor during its activation includes a high energy barrier (Sigg, Qian and Bezanilla, 1999). As has long been known, a high energy barrier converts the drift-diffusion of the particle into a movement that, under some conditions, may be approximated by the discrete state model, as formulated in the so-called Kramers’ approximation (Cooper, Gates and Eisenberg, 1988; Hänggi, Talkner and Borkovec, 1990; Eisenberg, Kłosek and Schuss, 1995)^1^. By contrast the drift-diffusion model predicts a variance independent of the mean current when the energy profile does not include a high barrier (Sigg, Qian and Bezanilla, 1999).

Thus, a simple system of a charge moving along an energy profile (without a significant barrier), does not produce shot noise. But a structurally more complex system that approximates a voltage sensor might have different properties, and different shot noise. Much structural and functional data show that during activation, gating charges move across two large vestibules separated by a short and highly hydrophobic gating pore impermeable to ions and water (Long *et al.*, 2007; Lacroix *et al.*, 2014). Several studies show that the electric field is negligible in the large vestibules, due to the high ion mobility there, and a significant voltage drop is only present inside the short hydrophobic gating pore (Islas and Sigworth, 2001; Asamoah *et al.*, 2003; Ahern and Horn, 2005). While the charged sensor is in the vestibules, its movement hardly produces gating current. On the other hand when the gating charge moves out of the vestibules and through the gating pore, gating current noise is produced with particular properties, as we will see in this paper.

We recently proposed a computer model of the voltage-dependent gating of Shaker channels where the voltage sensor is treated as a charged Brownian particle moving within a voltage sensor domain having geometrical and electrostatic properties taken from structural data. Notably, in this model the energy profile experienced by the voltage sensor during its movement is self-consistently evaluated using the laws of electrodynamics (Catacuzzeno, Sforna, Franciolini, and Eisenberg 2020; Catacuzzeno and Franciolini 2019; Catacuzzeno, Sforna, and Franciolini 2020). The model was shown to produce macroscopic gating currents quite similar to those experimentally observed.

In this paper we investigated in depth the properties of the gating current fluctuations produced by the model, and the main parameters modulating them. To do this, we first constructed a simplified model of the voltage sensor domain which differs from our full model (Catacuzzeno and Franciolini, 2019), and also differs from a realistic ion channel, because it does not include the permanent charges present near the voltage sensor domain. The simplified model also has a voltage sensor that moves over an energy profile that we choose arbitrarily, instead of being assessed self-consistently from all the charges present in the system. Although we are fully aware that this is a heavy simplification of the reality, this model allows us to connect with the existing biophysical literature, and to investigate how the gating current fluctuations vary with conditions. With the knowledge acquired with the simplified model, we will then calculate and study the fluctuations of gating current using a un-simplified and self-consistent model of voltage gating.

## The model

In this paper we deal with two different Brownian models of the voltage sensor, referred to as the “Simplified” and the “Full” model. In the Full model, already presented in the original paper (Catacuzzeno and Franciolini, 2019) and slightly improved in (Catacuzzeno, Sforna, Franciolini, and Eisenberg 2020), the geometry and charge distribution of the voltage sensor domain have been deduced from the structural atomic model of the Shaker K channel (Henrion *et al.*, 2012), and the voltage sensor dynamics are computed from the Langevin’s equation, while self consistently assessing the electrostatic potential that drives the voltage sensor dynamics. The self-consistent potential is computed from the Poisson equation, considering all the charges present in the model. To construct the Simplified model we started from the Full model, where we made several simplifications in the geometry and charge distribution that would allow us to better understand the qualitative properties of the gating current fluctuations and compare them to the existing literature. Most importantly, in this model we were able to arbitrarily choose the energetic profile driving the voltage sensor dynamic, a shortcoming that allowed us to simulate the dynamics of the voltage sensor under different energy profiles, essential to understand several key concepts at the origin of gating current fluctuations. Although the main equations used in the two models are the same, and have already been presented in the previous papers (Catacuzzeno and Franciolini, 2019), we will present them again in the context of the Simplified model, so that the reader can have a deep understanding of the simplification performed as compared to the Full model.

### Structure of the voltage sensor domain

In our model the VSD was approximated by an hourglass-shaped geometrical structure consisting of a water inaccessible cylindrical gating pore, having a length of 0.4 nm and a diameter of 1 nm, flanked by internal and external water accessible vestibules having a length of 3.1 nm each, and a conical shape opening with a half angle of 15° into two hemispherical subdomains of bath solution, both having radii of 1 μm. Each vestibule had a total volume of 7.9 nm^3^, thus allowing the simultaneous presence of only few ions at physiological conditions. This geometry allows the formulation of the model in one spatial dimension consisting in a main axis perpendicular to the membrane and passing through the gating pore. In the numerical simulation the main axis was divided into subdomains of constant step-size within the VSD, and a step-size increasing geometrically going outwards in the two bath solutions. Using this subdivision the surfaces separating adjacent sub-volumes (or slabs) were circles inside the gating pore, spherical caps in the vestibules, and hemispheres in the baths, each one contacting perpendicularly the channel wall (Figure 2A). The S4 segment does not occupy space in either vestibules, since it contributes to form the vestibule walls together with the other parts of the voltage sensor domain (S1-S3), as seen in the available crystal structure. The crystal structure shows that the extracellular vestibule is formed by a departure of the S3-S4 segments from the S1-S2 (Long *et al.*, 2007). We include the charges on the S4 segment as charge density in the volume grids of the model. The S4 charge profile (ZS4, expressed in e_0_ units) was built by considering 1 or 4 positive charges (depending on the simulation performed), each giving rise to a charge profile normally distributed with a standard deviation of 0.1 nm (in some cases we varied the standard deviation of the distribution). In our model the S4 segment was assumed to be a rigid body, and its position was represented by the variable xS4, expressing the distance of its midpoint from the center of the gating pore. During the simulation the voltage sensor was allowed to move through the gating pore and vestibules up to a maximal displacement xS4 of ±1.8 nm, by imposing fully reflective boundary conditions (i.e. if movement exceeds that displacement the particle is reflected inside the boundary by an amount equal to the exceeded displacement)^2^.

### Ion electro-diffusion

We assumed that the intracellular and extracellular walls of the VSD are bathed by ionic solutions containing 140 mM of positively and negatively charged monovalent ions, that can freely move in the baths and vestibules of the VSD, but cannot enter the gating pore, due to water and ions inaccessibility. Given the very small volume of the vestibules (about 7.9 nm^3^), a concentration of 140 mM there would correspond to a mean number of ions present inside close to one. The gating pore was assumed to have a relative dielectric constant (ε=4) much lower than in the bathing solution (ε=80): water and ions do not enter the gating pore. Ions were subjected to electro-diffusion governed by the following flux conservative equation:

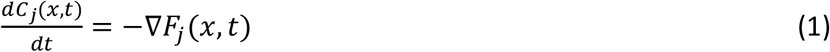

where *C_j_(x,t)* is the concentration of ion j, t is the time, ∇ is the spatial gradient operator, and *F_j_(x,t)* is the flux (mole per second per unit area) of ion j, given by the Nernst-Planck equation:

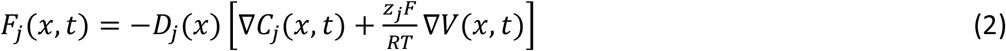

where *D_j_(x)* and are the diffusion coefficient profile and the valence of ion j, respectively, F, R and T have their usual meanings, and *V(x,t)* is the electrical voltage profile.

As also done in our previous model of voltage gating, we assumed steady-state for the dynamics of the electrolyte ions:

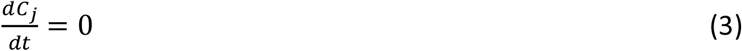

The validity of this approximation, which is based on the finding that ions relax on a timescale much faster than the movement of the S_4_ segment, has been fully demonstrated in (Catacuzzeno and Franciolini, 2019) and is a valid and necessary approximation in nearly all Langevin models (Eisenberg, Kłosek and Schuss, 1995).

The electrical voltage profile *V(x)* was assessed from the net charge density profile *ρ(x)*, using the Poisson’s equation

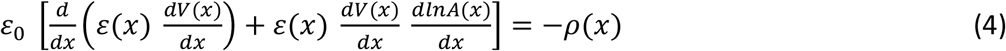

Where ε_0_ = 8.854⋅10^−12^ F⋅m^−1^ is the vacuum permittivity, *ε(x)* is the position dependent dielectric coefficient, *V(x)* is the electric potential and *A(x)* is the position dependent surface. The charge density profile was generated from the gating charges and the ions in solution:

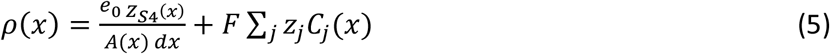

Eqns. (1) and (4) were iteratively solved until finding a steady-state solution for each allowed position of the voltage sensor, and the ion concentration profiles stored for the assessment of the gating currents (see later). The iteration made the treatment self-consistent, and always converged in our experience. In the supplementary data we report details of the numerical solution of the PNP system represented by eqns. (1) and (4).

### Movement of the S_4_ segment

In our model while ions where described by their concentration profiles (i.e., their mean number densities) (see above), the voltage sensor was treated explicitly as a moving particle occupying a well-defined position. More specifically, the S_4_ segment was assumed to move in one dimension as a charged Brownian particle, with dynamics governed by the following Langevin’s equation

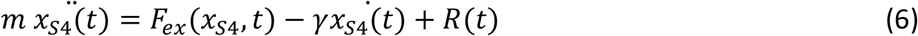

Here *x_S4_(t)* represents the position of the voltage sensor, *m* is the mass of the particle, *F_ex_(x_S4_,t)* is the external force acting on the particle, and R(t) is a random force due to the collision of the fluid and the rest of the protein on the S_4_ segment, which has a probability distribution with zero mean and second moment given by:

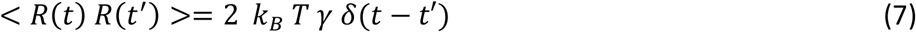

where *k_B_* is the Boltzmann constant, T is absolute temperature, and δ is the delta function. γ, the friction coefficient of the S_4_ voltage-sensor, was chosen to give, under the various conditions tested, gating current kinetics in reasonable agreement with the experimental data. More specifically the approach we used to set this parameter was to change it until the time course of the output gating currents was similar to that observed experimentally. For instance, when we varied the energy profile experienced by the particle, the time course of the gating currents varied accordingly (it was very sensitive to the height of the barrier). Thus we had to change γ in order to obtain gating currents with time courses similar to those observed experimentally.

In our gating model the friction experienced by the moving voltage sensor, γ, was assumed to be the same everywhere (and at any time, i.e., to be position- and time-independent). We considered this simplification reasonable even though the friction for the S4 segment is certainly very different inside the hydrophobic plug as compared to the solution. The reason for this assumption is that a portion of the S4 segment, of essentially constant length, is always inside the hydrophobic plug, and this part is arguably the one that mostly contributes to the overall friction of the moving sensor (as compared to the portions moving in the solutions outside the hydrophobic plug). We believe that this is true regardless of the specific residues on the S4 segment that are in the hydrophobic plug at different times during activation.

In the very high friction limit the inertial term 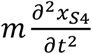 in eqn. (7) is much smaller that the friction term γ*x_s4_(t)*, and thus we arrive at the following stochastic differential equation (Titulaer, 1978, 1980; Miguel and Sancho, 1980; Eisenberg, Kłosek and Schuss, 1995):

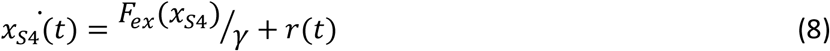

Where r(t) is a random Gaussian term with zero mean and second moment 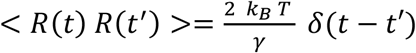. Eqn. (8) may be written in the form of the following stochastic differential equation, discretized with a Euler scheme (Higham, 2001):

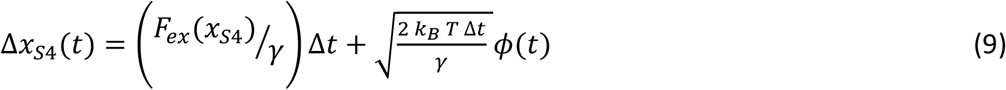

where *ϕ(t)* represents a normally distributed random variable with zero mean and unitary variance. Based on eqn. (9), the position of the particle may be found at each time-step Δt as 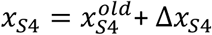, where 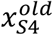 represents the initial position of the particle. The particle, as already stated, was allowed to freely move in the range *x_s4_* = ±1.8*nm*, by imposing elastic boundary conditions. More specifically we applied a totally reflecting boundary condition, using the following algorithm: if at a certain time-step the particle reaches a position x_1_>1.8 nm, its new position is set to 1.8-(x_1_-1.8), that is the particle is redirected inside the allowed spatial range by an amount exactly equal to the excess displacement. Similarly, if the particle in a certain time step reaches the position x_2_<−1.8 nm, then the new position of the particle will be −1.8 + (−1.8 − *x*_2_). In our model the external force acting on the S_4_ segment, *F_ex_ (x_S4_)*, is assessed as the negative gradient of the energy profile experienced by the voltage sensor, *G_tot_(x_S4_)*, which was arbitrarily chosen and varied during the study:

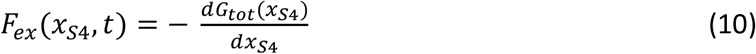

In our simulations of a single S_4_ segment, eqn. (9) was solved using a Euler scheme, with a time-step of 1 μs and using a normally distributed random number generator from (Press, 1992). We verified that the variance-mean current plot, resulting from thousands of simulations, as well as the properties of the microscopic current did not vary when further reducing the time-step.

### Assessment of gating current

As our main goal was to compare the output of the model with the experimental results, we computed the gating current exactly as it is normally done in experiments, that is, by assessing the ionic current measured at the intracellular and extracellular electrodes positioned far from the voltage sensor domain (I_gL_ and I_gR_, respectively). More specifically, the gating current was assessed by analyzing the net charge changes (with time) in the left (or alternatively in the right) bath, on the assumption that ions cannot pass through the gating pore, and by applying charge conservation equations:

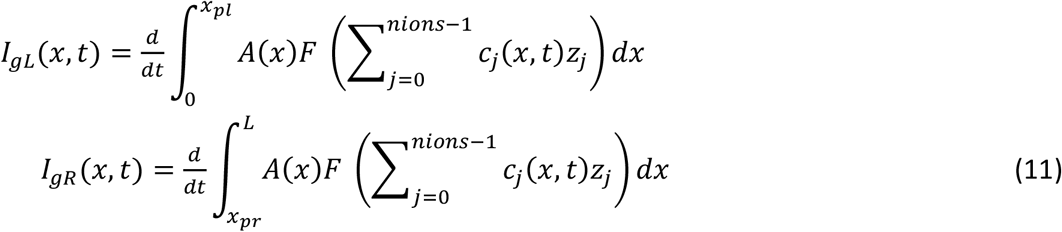

Where x_pl_ and x_pr_ are the left and right edges of the gating pore, F is the Faraday constant, and Zj and Cj(x,t) are the valence and the concentration of ion j. We verified in every situation that identical results were obtained for I_gL_ and I_gR_.

Maxwell’s equations guarantee that the total current in a series system is equal everywhere, at any time, no matter what the microphysics of conduction (Maxwell, 1965; Eisenberg, 2019; Eisenberg *et al.,* 2019). The total current is the [flow of ionic charge] + [the flow of gating charge] + [the displacement current], which is usually over approximated as *ε_o_ε_r_ ∂E/∂t.* We thus checked our model for the spatial conservation of current in the calculation of the microscopic gating current. As shown in Supplementary Figure 1, the model respects the spatial conservation of the total current throughout the entire domain, suggesting that the lack of a self-consistent treatment of the potential barrier is not injecting significant current into the system, as it might.

### Filtering of the current

The microscopic gating current is very noisy because of the high frequency Brownian movements of the voltage sensor. The noise obscured individual shot current events, and required the application of a digital filter. Since one of our goal was to compare our observation with experimental results, we usually used a digital filter reproducing the effect of an 8-pole Bessel filter with a cutoff frequency of 8 kHz, that is the same condition used in the experimental determination of the gating current fluctuations from Shaker K channels (Sigg, Stefani and Bezanilla, 1994). The C code for this filter was from (https://www-users.cs.york.ac.uk/~fisher/mkfilter/). Since our digital filter, as any Bessel filter, produces a characteristic ringing, in the detailed analysis of shot currents (Figure 5) we used a well-behaved Gaussian digital filter, which in addition allows the use of analytical expressions for its output (Colquhoun and Sigworth, 1995). In the implementation of the Gaussian filter, the output of each data point in the time course was:

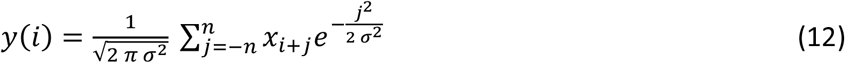

where *σ* = 0.1325/*f_c_,f_c_* is the cutoff frequency expressed in units of the sampling rate, *x_i+j_* of the unfiltered point at the (*i+j*)th position, and n was set to 70 to cover all the Gaussian coefficients significantly higher than zero.

### Overall algorithm for the Simple Model

In this section we report the overall algorithm used for the assessment of the time course of a microscopic gating current and of the variance vs mean current plots (Figure 1). At the beginning of the computation, the spatial profile of all the geometrical parameters (center position, surface, and volume of the slabs over the whole spatial domain) as well as of the various position-dependent parameters (dielectric constant, diffusion coefficients, charge density carried by the voltage sensor) were set. At this stage, for the Simplified model, we also set the energy profile experienced by the voltage sensor, instead of being computed later from the solution of the PNP system as in the case of the Full model. We then proceeded with the solution of the PNP system (eqns 1-5) for each possible position of the voltage sensor, using the numerical algorithm reported in the Supplementary data. Namely, the voltage sensor was positioned within each allowed volume slabs (those having a center position in the range of ±1.8 nm), and the ion concentrations and electric potential profiles at equilibrium were assessed and stored for later use. More specifically, the ion concentration profiles will be used for the assessment of the microscopic gating currents, whereas the electric potential profiles will be used in the Full model to assess the energy profile experienced by the voltage sensor, and driving its movement (Catacuzzeno and Franciolini, 2019a). Once the concentration and electric potential profiles for each possible position of the voltage sensor have been assessed, the time course of the voltage sensor position was assessed from eqns 8 and 9. More specifically, starting from an initial position, at each timestep Δt the position of the voltage sensor (x_s4_) is uploaded by X_s4(new)_=X_s4(old)_+Δx_s4_, where Δx_s4_ is assessed by eqn (9) using a random number generator to obtain *ϕ(t)*. During the computation of the time course of the voltage sensor position we also assessed the microscopic gating current using eqn 11 (to calculate the time derivative present in this eqn we used as information the overall ionic charge in the right (or left) bath at the current and previous time-steps). At the end of the simulation we filtered the gating current using either a Bessel or Gaussian filter as reported in the section “Filtering of the current”.

**Figure 1.**
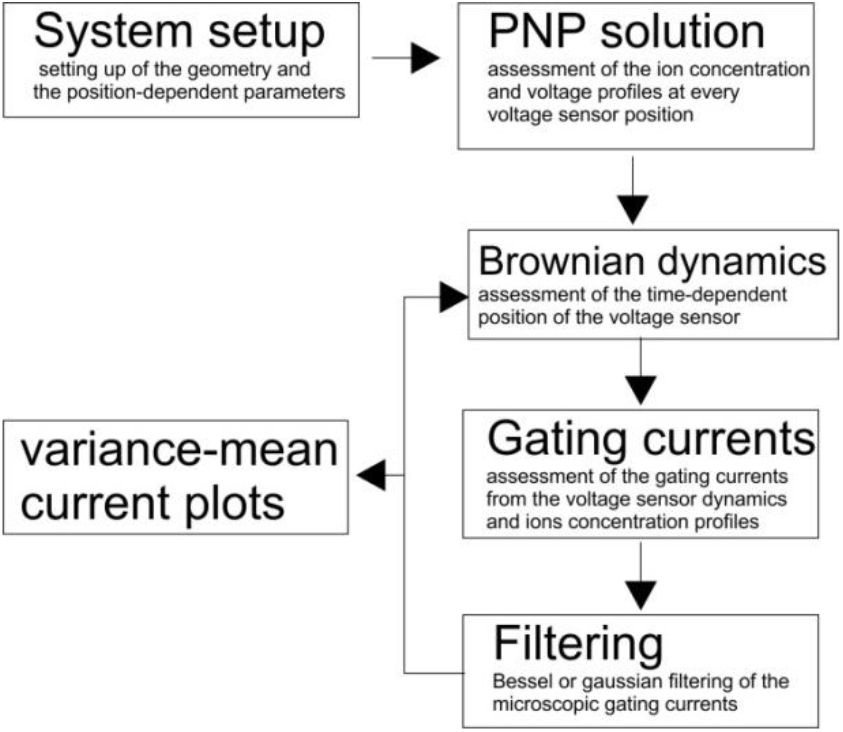
Scheme showing the various steps performed during the simulation of the microscopic gating currents and of the variance vs mean current plots. See text for details.

The variance vs mean current plot was made by repeating the simulation of the voltage sensor dynamics (eqn 8 and 9) tens of thousands of times, and at each time-step we assessed the mean and variance of the signal. In this routine there was no need to assess at each simulation the geometrical parameters, nor the voltage and concentration profiles associated to each voltage sensor position, since the conditions of the simulations were maintained constant.

## Results

### The Simplified model

The main properties of our Simplified model, derived from the Full model of (Catacuzzeno and Franciolini, 2019), are shown in Figure 2A. The voltage sensor domain consists of a short, cylindrical gating pore (5 Å radius and 4 Å length), with adjacent conical vestibules (3.1 nm long), and bath domains extending for 1 μm on both sides of the voltage sensor domain. Ions (monovalent, positive and negative, at a concentration of 140 mM) and water can freely move in the baths and vestibules, but cannot access and cross the gating pore. Further, the four charges on the voltage sensor were initially concentrated in one point, with a charge density normally distributed in space with a standard deviation of 1 Å. The four charges move back and forth across the gating pore and part of the adjacent vestibules (±1.8 nm, red arrow in Figure 2Ab), and this movement is modeled as a discretized Brownian stochastic process, and described using a discrete version of the Langevin equation of Brownian motion. Due to the very slow motion of the voltage sensor as compared to the ions in solution, we assumed that at each time steps anions and cations will instantaneously modify their concentration profiles in order to screen the gating charge (Catacuzzeno and Franciolini, 2019). More specifically the gating charge, together with the cations and anions in the baths, creates an electrostatic potential (Figure 2Ad) that in turn rearranges the ion concentration profiles in the bath (Figure 2Ac). Anions concentrate close to the gating charge (red lines) and cations deplete at the same location (black lines). When the gating charge moves, the ion concentration and the electric potential profiles will move accordingly, as shown by the two positions represented in Figure 2Ab-d (continuous and dash lines, respectively).

**Figure 2.**
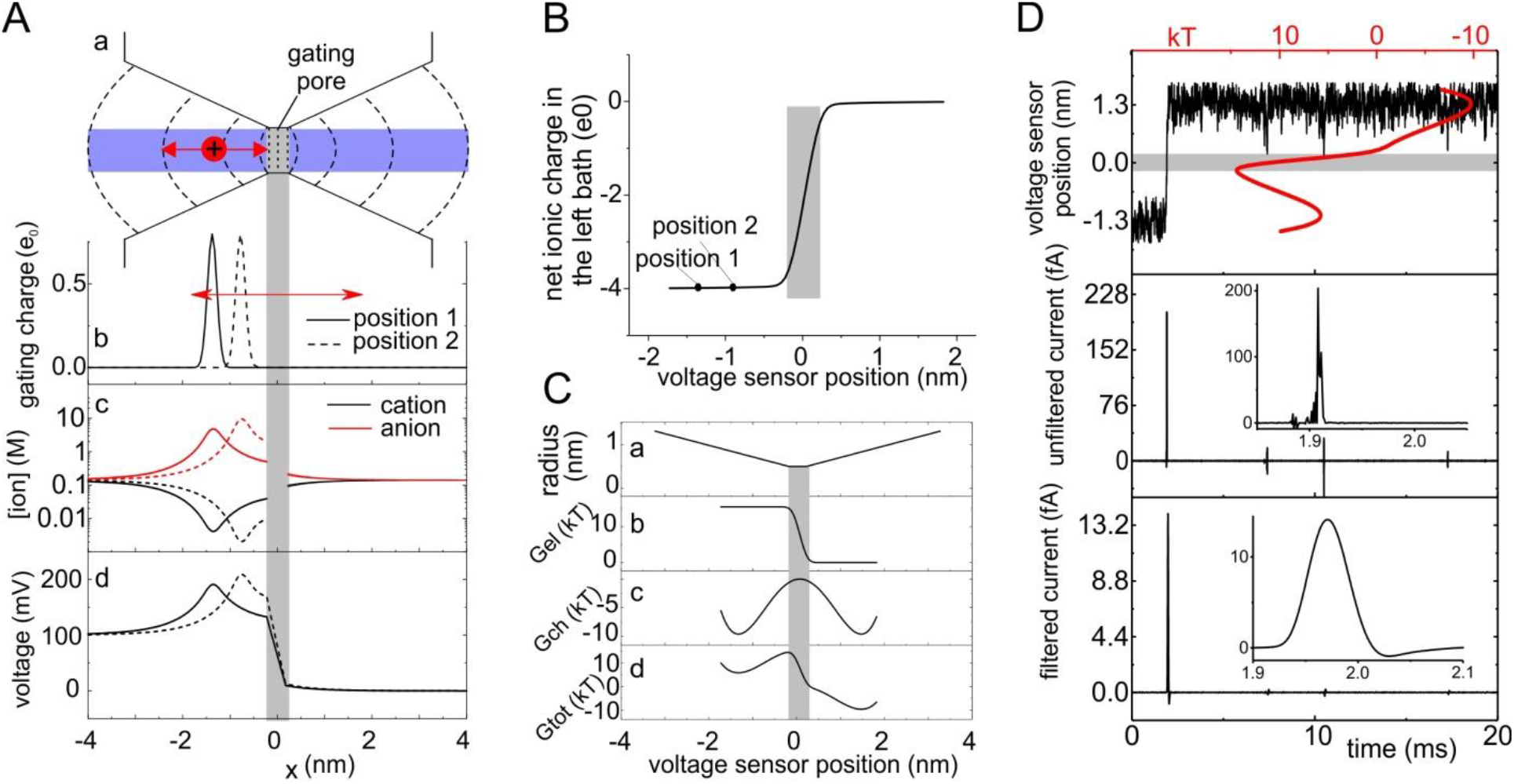
Properties and output of the Simplified model. **A)** Panel a shows a sketch of the voltage sensor domain considered in the Simplified model, with the gating pore and two adjacent vestibules. The dashed lines represent surfaces used to divide the space into sub-volumes. Panels b-d are plots of the position-dependent gating charge density, anion and cation concentrations and electrostatic potential, assessed at 0 mV of applied potential. The concentration and electrostatic potential profiles are shown for two different positions of the voltage sensor (solid and dashed lines, respectively). Notice that at the ion concentrations reached, only few ions will be present inside the vestibule at each time. **B)** Plot of the ionic net charge in the left compartment (bath and vestibule) as a function of the position of the voltage sensor. The two positions labeled “1” and “2” correspond to the two positions of the voltage sensor considered in panel A. **C)** Electrostatic, chemical, and total (electrostatic + chemical) energy profiles experienced by the voltage sensor during its movement through the gating pore. As stated in the text, in the Simplified model we arbitrarily choose the energetic profile experienced by the particle. This profile contains the overall energy charges experienced by the sensor, including the electrostatic voltage profile created by the voltage sensor and ions, and all other kinds of chemical and electrostatic interaction with the voltage sensor domain. **D)** Trial simulation performed with our Simplified model, using the energy profile shown in panel C and a friction coefficient of 2 10^−6^ Kg/s. From top to bottom the panel shows: the position of the voltage sensor (i.e., of the gating charge) from the center of the gating pore, the unfiltered gating current produced by the movement of the voltage sensor, and the same gating current filtered with a digital, 8-pole Bessel filter with a cutoff frequency of 8 kHz. The red line in the top panel represents the energy profile encountered by the voltage sensor. It is superimposed on the time course of the voltage sensor position to show that the voltage sensor spends most of its time in the energy wells present in the two vestibules (we used the same graphic style used by (Sigg, Qian and Bezanilla, 1999) to make comparisons with earlier work easier).

In our Simplified model the gating current recorded by the bath electrodes (located far from the gating pore) equals the charge (per unit time) that the electrodes need to put into the baths^3^, or absorb from them, to maintain electro-neutrality^4^. This means that a (gating) current is detected by the electrode only when the movement of the gating charge results in the addition or subtraction of charges into or from the baths. If during the time step Δt, the net charge in one bath changes by ΔQ, this means that the movement of the gating charge has added to (or subtracted from) the bath, and as result to the electrode, that same amount of charge, resulting in a recorded gating current of ΔQ/Δt. Figure 2B reports the net ion charge contained in the left bath as a function of the position of the voltage sensor. When the voltage sensor is in position 1, i.e. it is well inside the left bath, the net ionic charge there is −4e_0_ (to compensate for the +4e_0_ of the gating charge, and keep electro-neutrality in the bath). Slightly moving the voltage sensor to the right, for instance from position 1 to position 2 in Figure 2Ab, we see that the net ionic charge remains close to −4e_0_, as expected given that the position of the voltage sensor remains well inside the left bath.

In other words, this movement does not generate any gating current because the gating charge has not entered the gating pore and thus has not subtracted charges from the left bath. Only when the voltage sensor enters inside the gating pore (gray region) a sensible change in the net charge of the left bath is detected, and this originates a gating current.

Finally, in our Simplified models the voltage sensor moves in an energy profile that we choose arbitrarily. Although we are aware that any physical model of voltage-dependent gating should self-consistently compute the energy profile, as we did in (Catacuzzeno and Franciolini, 2019), this shortcut turned out to be decisive in helping us understand how the gating current depends on the shape of the energy profile and on many other parameters. Later we will show results where this approximation is removed. As shown in Figure 2C, the energy profiles experienced by the gating charge can be thought of as the sum of an electric component (G_el_), produced by an imposed membrane potential entirely and linearly dropping within the gating pore (Figure 2Cb), and a “chemical component” (G_ch_) shaped as two wells and one barrier, with the energy barrier centered in the middle of the hydrophobic gating pore, and the wells located in the baths, where the gating charges are balanced by counter-charges (Figure 2Cc)^5^. Figure 2Cd shows the overall energy profile experienced by the gating charge, resulting from the addition of the electric and chemical components. Given the symmetry of the chemical component in the Simplified models, and resulting identical intracellular end extracellular energy minima, at zero applied potential the voltage sensor does not experience a net force in either direction, and thus it is expected to spend equal time in the left and in the right vestibule.

Figure 2D shows a typical outcome of the simulation of a microscopic gating current, obtained with an energy profile that includes a high barrier (>5 kT) across the gating pore, in addition to a voltage drop of 100 mV (cf. Figure 2C). The top panel shows the position of the gating charge with respect to the gating pore, as function of time. Superimposed and in red the same plot also reports the total energy profile encountered by the voltage sensor (same as in Figure 2Cd). Because of the presence of a high barrier separating the two wells, the gating charge is almost always in one well or the other, except for the few tens of microseconds during which the gating charge is crossing the gating pore (or the barrier). The central panel also shows that the fluctuating movement of the charge within the vestibules produces a negligible amount of current. By contrast the passage of the gating charge across the gating pore causes a needle-like current (expanded on the time domain in the inset). The features of this current spike, that coincides with the gating charge crossing the gating pore, recall the shot current postulated to occur when the voltage sensor crosses the membrane (Conti and Stühmer, 1989; Sigg, Stefani and Bezanilla, 1994). The bottom panel displays the same current trace after being filtered at 8 kHz with an 8-pole digital Bessel filter.

#### Constructing variance-mean current plots and recovering the apparent gating charge

The variance-mean current plots have been classically used to assess the apparent gating charge, q_app_, that moves across the gating pore during channel activation. The mean current and the corresponding variance with our Simplified model were obtained from the ensemble of thousands of filtered microscopic gating currents, and used to construct the variance-mean current plot (Figure 3), which shows a clear linear dependence between the two variables.

**Figure 3.**
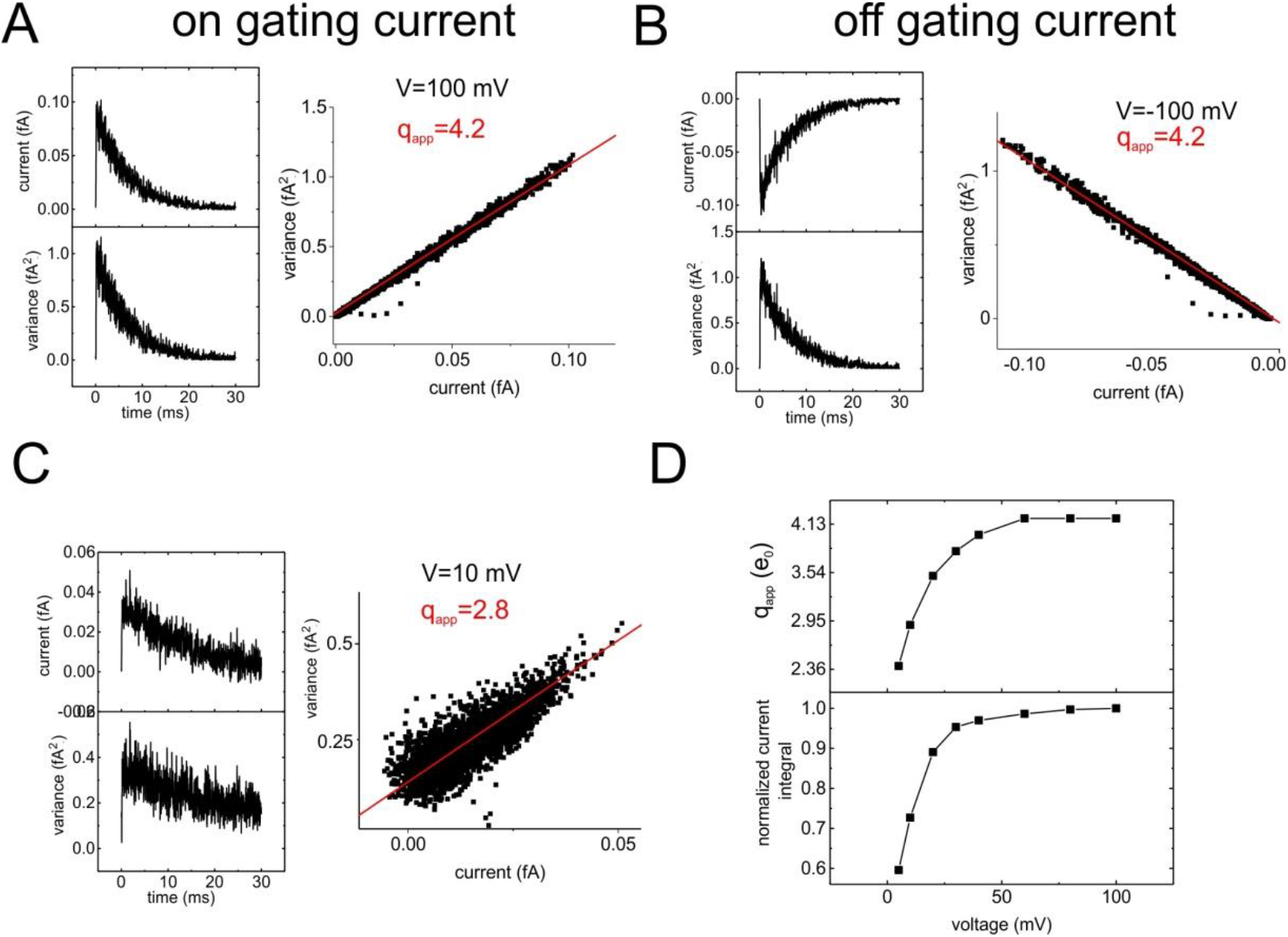
Simulations of variance vs mean current plots with the Simplified model. **A)-C)** The left panels show the time course of the mean (top) and variance (bottom) assessed from 10,000 simulated microscopic gating currents, while the right panels represent the variance vs mean plot obtained from the same data. The red lines represent the fit of the data with eqn. (13) to which a constant, current-independent variance was added (Sigg, Qian and Bezanilla, 1999), giving the reported q_app_. Panels A and C were obtained with the voltage sensor initially placed at the most intracellular position (−1.67 nm), and stepping the applied voltage to either +100 or +10 mV. They thus represent ON gating currents produced by the forward movement of the voltage sensor. By contrast in Panel B the voltage sensor started from an outward position (+1.67 nm) and the applied voltage was maintained at −100 mV (OFF gating currents). **D)** Plot shows q_app_ and normalized current integral (that is, a measure of the activation degree) as a function of the applied voltage. While at voltages that ensure an essentially irreversible activation q_app_ results close to the real charge carried by the voltage sensor, at lower voltages this parameter tends to be sensibly lower.

To be connected to the widely used literature, the data were fitted using the following equation, valid for a two state Markov model, and at applied potentials where the activation process becomes essentially irreversible:

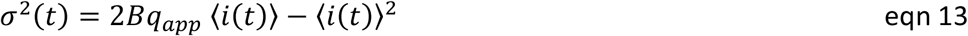

where B is the cutoff frequency of the filter, q_app_ is the apparent gating charge^6^, *σ^2^(t)* is the variance, and *〈i(t)〉* is the mean current. In the case considered in Figure 3A and B, i.e. ON or OFF gating currents obtained in the presence of a relatively high energy barrier, and at a potential where the process can be considered essentially irreversible, the fit gave an apparent gating charge q_app_, for both the ON and the OFF gating currents, of 4.2e_0_, close to the real gating charge considered in the model^7^. By contrast, at lower voltages that activate the voltage sensor only partially (consider that the activation V**1/2** in the model is 0 mV), the resulting q_app_ was sensibly lower (Figure 3C), in accordance with the limit that eqn 13 holds only for irreversible processes.

#### Dependence of the fluctuation properties on the voltage-sensor energy profile

We then repeated this type of analysis using always the same applied potential of 100 mV, but an energy barrier of lower amplitude^8^, to verify whether also in our model a q_app_ close to the actual gating charge is only obtained in presence of a high energy barrier. A q_app_ close to 4 was still obtained reducing the barrier height to 5 kT (Figure 4A). However, as shown in Figure 4B, further reducing the barrier height, while keeping the same voltage drop, the slope of the variance-mean current plot decreased (to give q_app_=3.6e_0_ for a barrier height of 2kT), as expected for the particle dynamics as it lost its high barrier, Markovian character. We finally removed the barrier altogether (0 kT). Under these conditions we expected to find a q_app_ close to zero from the variance-mean current plot, given the results of (Sigg, Qian and Bezanilla, 1999) that energy profiles without appreciable barriers generate time-(or current-) invariant noise (in the case of a position independent friction coefficient, as we have in our model). Notably, a significant slope of the variance-mean current plot (with a q_app_=2.6e_0_) remained even after the energy barrier within the gating pore had been set to zero (Figure 4C), suggesting that a shot current is present even under non-Markovian conditions, i.e. in the absence of well-defined discrete states for the voltage sensor. This conclusion seems to be supported by the simulations reported in Figure 4D where the microscopic gating current is shown for a model not including an energy barrier. Panels **a** and **b** of Figure 4D show that while the gating charge roams in the bath where no voltage gradient is present, no gating current is generated (see lower panels). A clear and significant current flickering is instead observed as soon as the gating charge enters and swiftly and unidirectionally traverses the gating pore, under the drive of the voltage gradient present. This rapid crossing of the gating pore produces a current resembling the shot observed in presence of an energy barrier, as clearly shown at the bottom of panel Db. Figure 4Dc reports representative traces showing the microscopic gating currents/shot events generated as the gating charge crosses the gating pore, in seven different trials.

**Figure 4.**
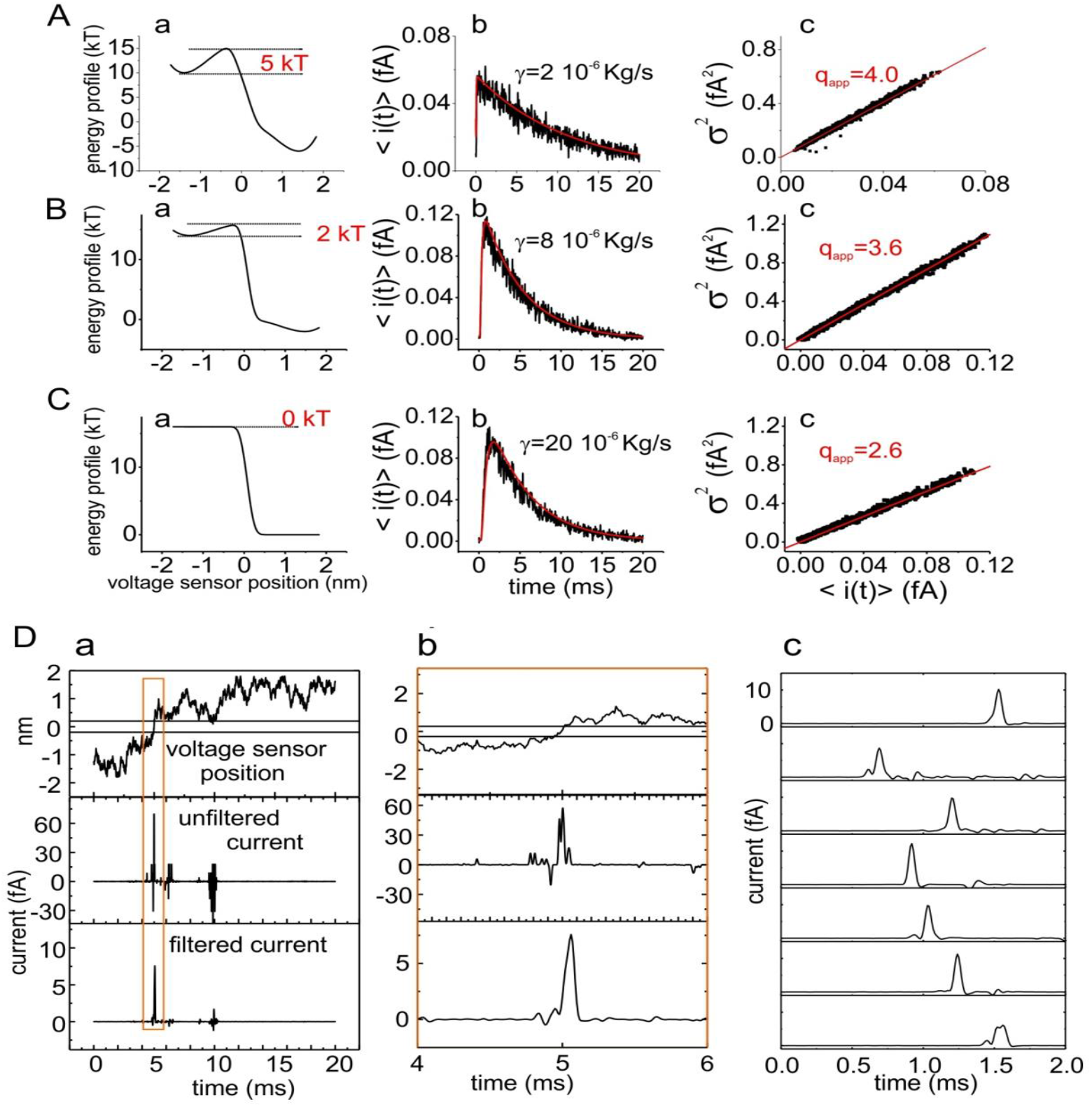
Presence of a shot noise in the absence of an energy barrier. **A)-C)** Assessment of the variance-mean current plot for models including a double-well energy profile characterized by different barrier heights, with the Simplified_**CC**_ model. Panels **a** report the energy profiles encountered by the gating charge during its movement across the gating pore. Panels **b** report the mean current resulting from ten thousands simulations of the microscopic current. Each simulated current was filtered with an 8-pole Bessel filter with a cutoff frequency of 8 kHz. Panels **c** show the variance-mean current plots. The red lines are the best fits of the row data with eqn. 1, resulting from a one-step Markov model. **D)** Typical outcome of simulations in the absence of an energy barrier (cf panel Ca). Panel **a** shows the gating charge position (top), unfiltered current (middle), and filtered microscopic gating current (8-pole Bessel, f_c_=8 kHz; bottom) for a typical simulation. Panel **b** reports a time expansion of the same simulation shown in a. Panel c reports the filtered microscopic currents obtained in 7 different simulations performed under the same conditions of panels **a** and **b.**

As a first attempt to analyze in detail the shape of the shot events in the absence of an energy barrier, using the simplified model, we tried to increase the cutoff frequency of the filter. As shown in the Supplementary Figure 2 both the amplitude and duration of the unitary events (shots) were strongly influenced by the filter cutoff frequency included in our model, suggesting a very fast underlying signal that has been significantly smoothed by our low pass filter. However, the increase in the filter cutoff frequency also resulted in the appearance of a high amplitude noise due to the currents generated from the high frequency Brownian motion of the gating charge, that obscured the shot current. We thus tried to deduce the shape of the signal underlying the filtered shot, by taking into account the effect of the filter. For the purpose of this analysis we filtered the microscopic current with a Gaussian filter (f_c_=8 kHz), since it is better-behaved and avoids most of the ringing usually observed with Bessel and other filters available for analog signals. As shown in Figure 5A, the filtered signal (black line) deviates significantly from the prediction of an instantaneous shot of charge filtered with a Gaussian filter (blue line), suggesting that the Markov assumption of an instantaneous movement of the gating charge across the gating pore is not appropriate, as expected in a model including no barrier in its energy profile. We also found that the shots could be well fit by an equation predicting the observed time course of a filtered current step (Figure 5A, red line) (Colquhoun and Sigworth, 1995), resulting in a mean amplitude and duration of the underlying current step of 17.6±2.1 fA and 35.9±5.3 μs, respectively (Figure 5B).

**Figure 5.**
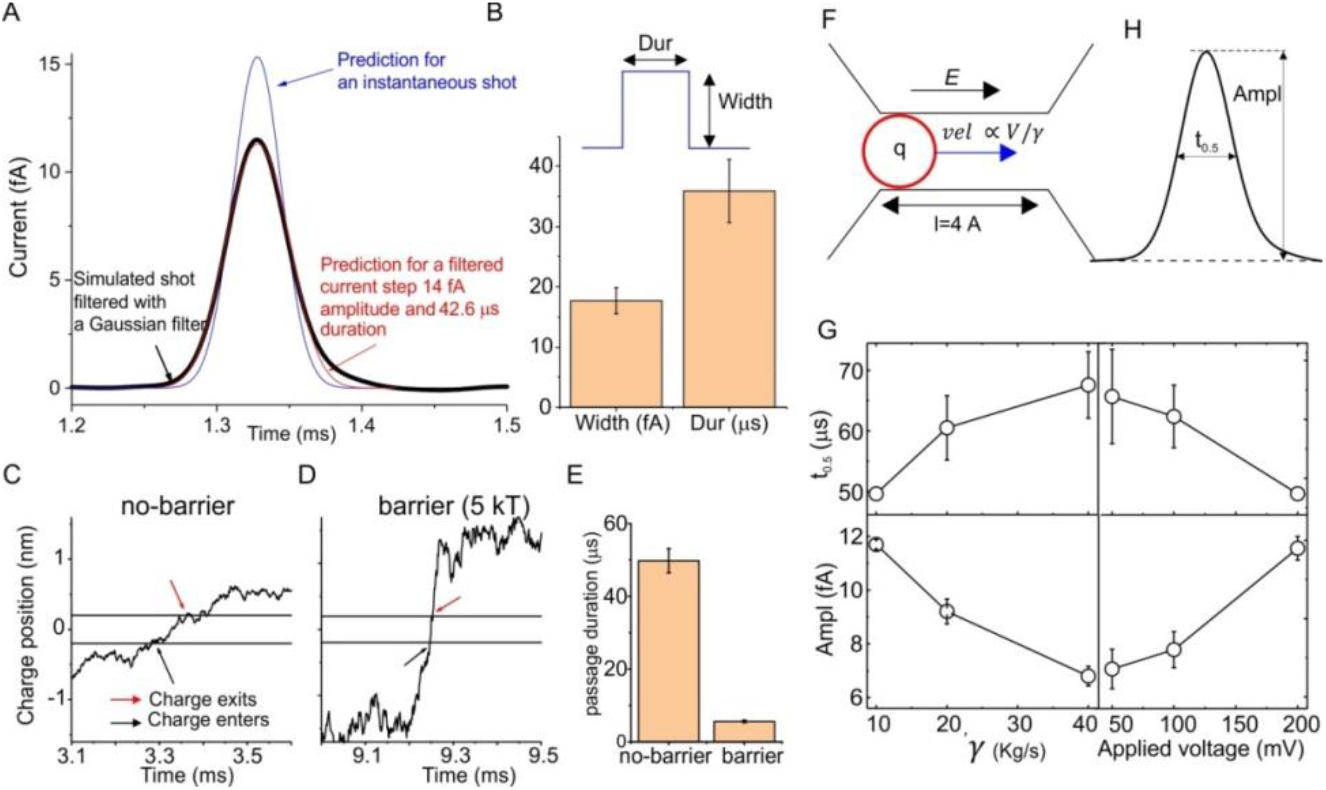
Analysis of the shot current observed in absence of energy barrier. **A)** The plot shows a typical shot current observed under no-barrier conditions (black line), filtered with a Gaussian filter with B=8 kHz. The blue line represents the response predicted for an instantaneous shot of charge filtered by a Gaussian filter, namely *i(t)* = *q h(t)*, where q is the gating charge and h(t) is the Gaussian response function, reported in the Appendix of the Supplementary data. The red line is a fit of a squared shot current of duration *Dur=42.6* μs and amplitude *Width=14* fA with the following equation for Gaussian filtered current steps from (Colquhoun and Sigworth, 1995): 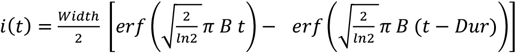. **B)** Plot of the mean Width and Dur parameters obtained by fitting 10 shot currents as shown in panel A, with the equation for a filtered current step. **C)** Time-dependent position of the voltage sensor obtained from a simulation with the Simplified model, with 100 mV of applied potential and no energy barrier. The black and red arrows indicate the time of entrance and exit of the voltage sensor into (from) the gating pore. **D)** Same plot as in panel C, except for the 5 kT energy barrier (same condition as in Figure 2A). **E**) Plot showing the mean time taken by the voltage sensor to pass across the gating pore, in the no-barrier and 5 kT barrier cases. **F)** Proposed physical mechanism of the generation of a shot current in our Simplified model with an applied voltage and with no energy barrier. Due to the presence of an electric field inside the gating pore, the gating charge is subjected to an electric force, thus generating a current. This current is predicted to be directly proportional to the applied voltage, and inversely proportional to the friction coefficient experienced by the voltage sensor during its movement. **H**) Illustration of a shot current and the parameters used to quantify it, namely the amplitude (Ampl) and duration at half amplitude (t_0_._5_). **G)** Plot showing the parameters Ampl and t_0_._5_ of shot currents, defined in panel H, as a function of the friction coefficient and of the applied potential.

We also observed that under the no-barrier condition the voltage sensor required a relatively long time to cross the gating pore, as measured by directly looking at the time course of the position of the voltage sensor (Figure 5C and E). This time was however much shorter in the model with a high energy barrier (Figure 5D and E, same energy profile of Figure 4Aa). To put it crudely, the voltage sensor took much longer to diffuse across the gating pore than to jump across a barrier. The drawing in Figure 5F sketches a possible physical mechanism for the shots observed in absence of an energy barrier. As shown earlier, the gating charge will generate a current only when it enters and moves inside the gating pore, where it senses the electric field. Here, it will experience an electrical force given by *F=qV/l,* where *q* is the gating charge, *V* the applied voltage and *l* the length of the gating pore. This force will remain constant for the entire length of the gating pore, and will drive the particle towards the right vestibule at a constant drift velocity *vel=F/γ=qV/(l γ).* The mean time taken to cross the gating pore will be given by 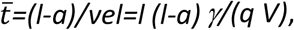 where the parameter *a* takes into account the fact that in our model the charge is not a point charge, and it needs to be a good way inside the gating pore ir to be experience most of the electric field (similar reasoning when the charge exits from the right boundary of the gating pore)^9^. Notably, as the charge moves inside the gating pore at a constant drift velocity, for the averaged time 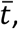 it will produce a constant current 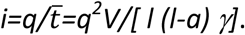 Using parameters *V*=100 mV, *l*=4 Å, *q*=4e_o_, *γ*=20e^−6^ Kg/s, *a*=0.5 Å we obtain 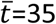 μs and *i*=18.2 fA, values reasonably in accordance with those obtained from fitting the signal to a filtered current step (Figure 5B). This simple treatment shows that duration and amplitude of shot currents observed in absence of an energy barrier depend on *V* and *γ*, as the analysis of the unitary events confirmed (Figure 5G).

We also looked for V- and γ-dependent shot noise by performing noise analysis. As shown in the Supplementary Figure 3, we did find a marked dependence of noise fluctuations on both parameters (in the absence of energy barriers), as assessed from the slope of the variance-mean current plot. Under these conditions the slope of the variance-mean current plot displayed a clear dependence on the friction coefficient (with a 4-fold change in γ resulting in a 2-fold change in the slope), and on voltage (increasing about 50% on doubling the voltage step). By contrast, no dependence on both parameters was found in presence of a high energy barrier, as expected from eqn. 13.

#### Analysis of the fluctuation properties with distributed charges on the voltage sensor

It is now well established that the first four arginines, positioned every third residue on the S4 segment, are the main carriers of the gating charge. These residues are separated by an α-carbon distance of 4.5-6.0 Å, depending on the assumed secondary structure for the S4 segment (α helix or 3.10 helix). We thus verified the consequences of a more realistic situation, with 4 unitary gating charges evenly distributed along the voltage sensor, each spread according to a Gaussian distribution with a standard deviation of 1 Å.

#### Dependence of the fluctuations on the inter-charge distance and energy barrier

As before, we assumed a linear voltage drop within the gating pore, resulting in an electric component of the energy profile with a staircase shape (Figure 6B). In addition, four energy barriers were added, associated with the passage of each unitary charge through the gating pore (Figure 6B). Figure 6A shows three different simulations, all performed with the same energy profile (Gtot) shown in Figure 6B, but using three different inter-charge distances, ranging from 4.5 to 8 Å. For a distance of 8 Å the simulations resulted in a q_app_ close to 1e_0_, indicating that under these conditions the passage of each charge through the gating pore may be viewed as a Markovian step carrying exactly 1 elementary charge^10^.

**Figure 6.**
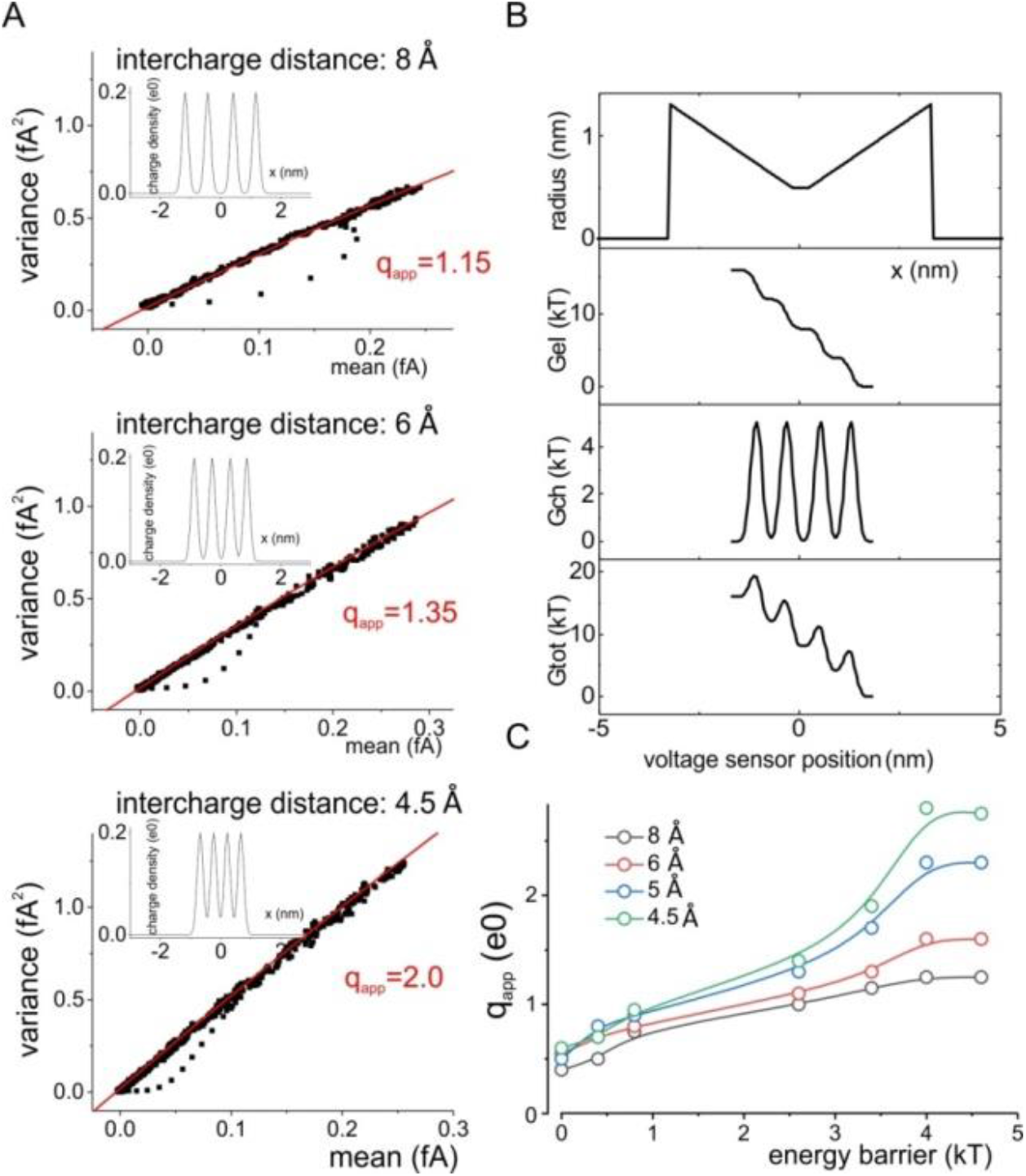
Gating current fluctuations with multiple gating charges on the voltage sensor. **A)** Variance-mean current plots obtained in simulations performed with the Simplified model, with four gating charges on the voltage sensor (charge density was normally distributed with a standard deviation of 1 Å). The three plots correspond to a different inter-charge distance (indicated). The insets in each plot show the corresponding gating charge distribution. **B)** Plots of the radius, electrical, chemical and total energy profiles for the simulation, shown in panel A, with an inter-charge distance of 8 Å. **C)** Plot of the apparent charge estimated from simulations performed at different inter-charge distances and barrier heights.

However, as the inter-charge distance gets smaller, and more similar to a realistic distance between charges located on an α helix, q_app_ increases, deviating significantly from 1e_0_. In Figure 6C we report the results of simulations performed at various inter-charge distances, and using varying heights of the energy barriers. Notable, as already seen with a single charge (cf. Figure 4), the q_app_ decreases while reducing the energy barrier (regardless of the inter-charge distance), but it never gets any close to zero, even in the complete absence of a barrier.

The increase in the apparent charge for low inter-charge distances might be caused by the fact that when the charges get closer the charge density distributions of two consecutive charges may partially be found simultaneously inside the gating pore. As expected on the basis of this interpretation, the q_app_ was found to be related to the spreading of the gating charge distribution. Increasing σ, the standard deviation of the Gaussian charge distribution, results in an increase in q_app_, while decreasing σ, and thus decreasing the charge overlap, results in a decrease in q_app_ (See Supplementary Figure 4).

#### Features of elementary shot events produced by distributed charges

We also studied in detail the shot events produced by models having two different inter-charge distances (8 and 4.5 Å), in order to find a mechanistic interpretation for the increased slope of the variance-mean current plot at shorter inter-charge distances^11^. As shown in Figure 7A-C, at inter-charge distance of 8 Å the responses produced by the passage of the voltage sensor through the gating pore most often (15 out of 20 simulations) produced four clearly distinct current peaks (see the four topmost responses in Figure 7B). In four simulations we found three peaks (see bottom response in Figure 7B), and in one we found only two peaks. As clearly shown in the running integral of the microscopic current (Figure 7C), in the 4-peak responses each peak corresponds to the passage of about one unitary charge through the gating pore. In contrast, in the four responses with three peaks we found that one of the peaks carried 2 unitary charges. In the one simulation with two peaks, one peak carried one charge, and three charges the other (not shown). These different responses are also evident in the charge amplitude histogram shown in Figure 7D, showing that most of the current peaks we detected under these conditions carry a charge of about 1e_0_. These results show that with the gating charges positioned relatively far one another, each charge goes through the gating pore more or less independently from the others, thus producing shot events carrying mostly one elementary charge. It is thus not surprising that in this condition the variance-mean current plot gives a q_app_ of about 1e_0_ (cf Figure 6A, upper plot), as predicted by a Markovian model with a 4-step activation process that should represent the sequential passage of the four gating charges through the pore.

**Figure 7.**
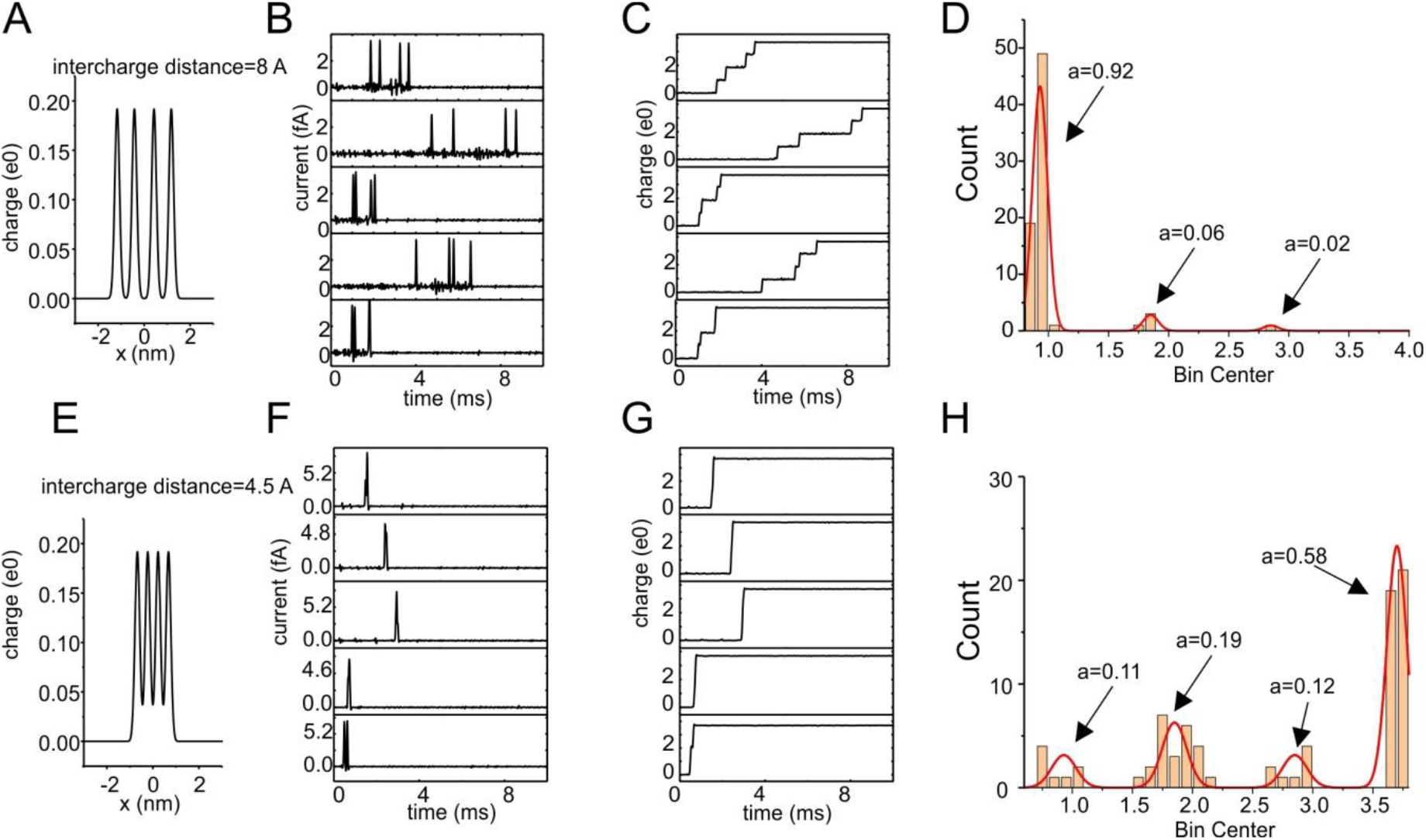
Evidence for multi-charge steps at low inter-charge distance. **A)** and **E)** Charge distributions for two Simplified models having a different inter-charge distance. In both cases an applied potential of 100 mV and an energy profile with barriers of 4.5 kT were considered. **B)** and **F)** Filtered current responses for the two models, showing that while at a relatively high inter-charge distance multiple peaks are clearly visible in the response, at lower inter-charge distance often the passage of the four gating charges occurs in only one step. **C)** and **G)** Running integrals for the responses shown in panels B) and F). **D)** and **H)** Histograms of the charge carried by the current peaks, accumulated from 20 (D) or 60 (H) responses as those shown in panels B and F. It is evident that for large inter-charge distances most of the current peaks carry a single gating charge, whereas for lower inter-charge distance peaks carrying multiple gating charges are significantly represented.

When the inter-charge distance is reduced to 4.5 Å, the analysis of the shot events gives a completely different scenario (Figure 7E-H) as one might expect since electrostatic and steric interactions are very hard to avoid at these close distances. In this case, in most of the simulations (40 out of 60) we only see one current peak, corresponding to the passage of a charge of about 4e_0_ through the gating pore in only one step. There were however several instances (20 out of 60) where the voltage sensor moved in two steps, each carrying 1, 2 or 3 unitary charges (see one such outcome in the bottom trial in Figure 7F and G). This suggests that when the gating charges are relatively close, they can pass through the gating pore together^12^, in various groupings, producing shot signals with integrals larger than one elementary charge. In this case it appears obvious that the variance-mean current plot will have a q_app_ higher than 1e_0_, since most of the shots (ca. 90% under these conditions; cf Figure 7H) carry a charge higher than 1e_0_.

The results presented above suggest that a q_app_ higher than 1e_0_ describes gating charges passing together through the gating pore. In order to clarify this point, we made a mathematical derivation for the predicted slope of the variance-mean current plot for shot noise events carrying a different number of unitary charges. The analysis assumes that the shot events are instantaneous, as in a true Markovian process, and thus the resulting predictions are valid only for very high energy barriers. As shown in Supplementary data the variance is predicted to be linearly related to the mean current, and its slope given by 2Bq_app_, where q_app_ is defined by the relationship:

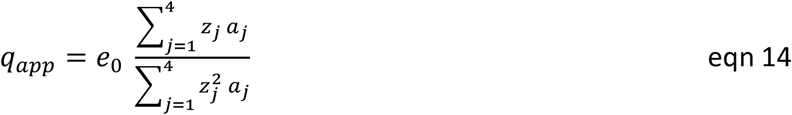

Here a_j_ is the frequency of occurrence of shots carrying z_j_ unitary charges. In the case of inter-charge distances of 8 Å and 4.5 Å, eqn. 14 results respectively in q_app_ of 1.13 and 3.3 (taking the mean charges and frequency of occurrences from the histograms in Figure 7D and H). These values are not dissimilar from those found by fitting the corresponding variance-mean current plots (cf Figure 6C).

#### Conclusions from the Simplified model

The results reported above suggest that caution should be used when interpreting the slope of the variance-mean current plot using the classic relationships of Markov large barrier theory (slope=2Bq, where q is the overall gating charge for a single step process, or the charge carried by the highest charge-carrying single step for a multi-step process). In fact:

1. the slope of the variance-mean current curve may be substantially lower than the value of 2Bq expected from Markovian theory due to the presence of not sufficiently high energy barriers (Figures 4 and 6C).
2. the slope of the variance-mean current curve may be substantially higher than zero even in the complete absence of energy barriers, i.e., when stable Markovian states cannot be defined (Figure 4, 5 and 6C).
3. In a multi-step process, as likely occurs in real channel gating, the slope of the variance-mean current curve may result substantially higher than the maximum charge carried by a single step, due to multiple charges crossing the gating pore together (Figures 6 and 7).

The simple model considered here is a reduction of the complete model and is probably not a unique reduction, as is so often in the case in approximations (of this perturbation type) in applied mathematics. Other approximations may exist and have advantages, but one of the main points of this paper is to point out the advantages of the Full model, with the obvious implication that it be used wherever practical. Given the history of computation, it is not unreasonable to suggest that what seems impractical today will be come routine in a few years.

### The Full model

The results presented so far were obtained with a simplified model of voltage gating, where no fixed charges on the voltage sensor domain were considered, the gating pore had a uniform dielectric constant, and most notably the energy profile for the voltage sensor movement had an arbitrary shape, not assessed self-consistently. With the knowledge acquired with the simplified model, we are now ready to evaluate the gating current fluctuations produced by our full model of voltage gating (Catacuzzeno, Sforna, Franciolini, and. Eisenberg 2020; Catacuzzeno and Franciolini 2019). This will allow us to verify how well the model predicts the experimentally observed variance-mean current relationship, and provide clues to understand its origin.

Figure 8 is meant to show that the full model is consistent with experimental data. Panels A and B show the variance-mean current plot obtained with the full model by pulsing to 0 mV, assessing the microscopic gating currents resulting from the motion of the voltage sensor, and filtering the response at a cutoff frequency of 8 kHz. When the variance-mean current plot derived from the gating current decay was fitted with the equation resulting from the Markov model (eqn. 13), we obtained a q_app_ of about 2.0e_0_. This value was not much different from the 2.4e_0_ obtained experimentally on Shaker channels (Sigg, Stefani and Bezanilla, 1994).

**Figure 8.**
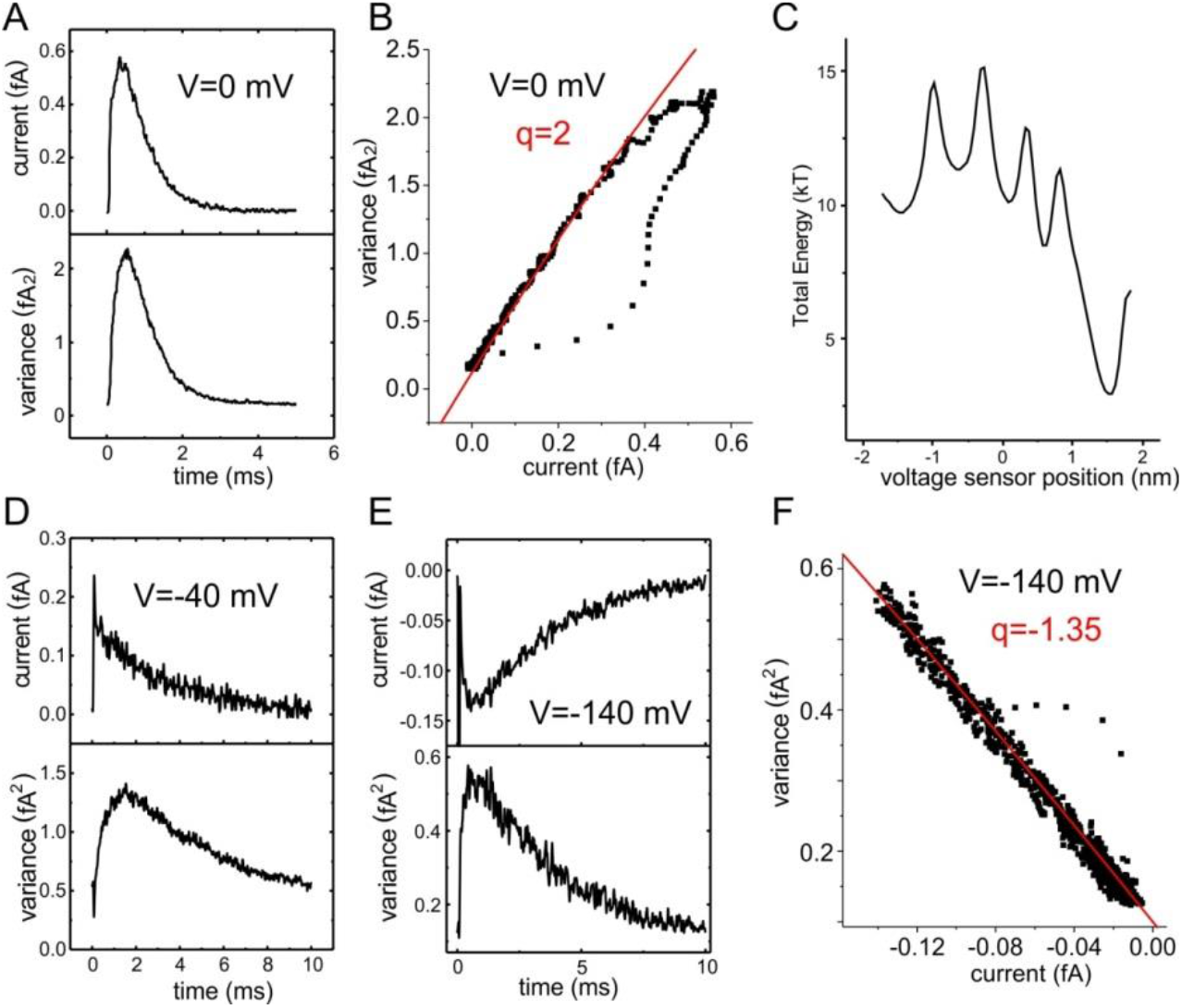
Gating current fluctuations in the Full model. **A)** Plots of the mean current and variance assessed from 10,000 simulated microscopic gating currents obtained in response to a depolarizing step to 0 mV (initial position of the voltage sensor −1.67 nm). Each simulated microscopic current was filtered with an 8-pole Bessel filter with a cutoff frequency of 8 kHz. **B)** Plot of variance vs mean current made with the data shown in panel A. The solid line represents the fit of the data from the decaying part of the gating current with eqn 13, with q=2.0 e_0_. **C)** Total energy profile experienced by the voltage sensor during its movement. Notice that this energy profile, differently from those arbitrarily chosen in the Simplified model, is self-consistently assessed with the Poisson equation, considering the effect of all the charges present in the system. **D) and E)** Plots of the mean current and variance assessed from 10,000 simulated microscopic gating currents obtained in response to a depolarizing step to −40 mV (D, initial position of the voltage sensor −1.67 nm), or to a repolarizing step to −140 mV (E, initial position of the voltage sensor 1.67 nm). **F)** Plot of variance vs mean current made with the data shown in panel E. The solid line represents the fit of the data from the decaying part of the gating current with eqn 13, with q=1.35 e_0_.

Nonetheless, when we saw the energy profile encountered by the voltage sensor during its movement through the gating pore (Figure 8C)^13^, we were initially rather surprised with this result. As already discussed in (Catacuzzeno and Franciolini, 2019), the profile displays 5 energy wells corresponding to the five stable positions of the voltage sensor (S4) while it moves through the VSD. The four energy barriers arise from the movement of one of the four gating charges inside the gating pore, where they experience destabilizing electric forces because of the low dielectric constant and the absence of counter ions inside the pore. Without the results obtained from our Simplified model, this energy profile could have suggested that a four-step Markov model, with each step carrying one charge through the gating pore, might be adequate to describe the activation process, as previously proposed (Zagotta, Hoshi and Aldrich, 1994; Schoppa and Sigworth, 1998). However, when we tested it with our Full Brownian model we obtained a q_app_ of 2e_0_, a value twice as large as the 1 electronic charge expected from a 4-step Markov model.

Figure 8D shows the mean current and variance time courses obtained for a depolarizing step to - 40 mV. A marked delay in reaching the variance peak, compared to current peak, is observed under these conditions, a result also obtained in experiments. Notably, when the noise analysis was performed for a hyperpolarizing step to −140 mV, starting with the voltage sensor in the activated position, we obtained a qapp value much lower than that resulting on depolarization to 0 mV (1.35 vs 2.0, Figure 8E and F). This result is not in accordance with experiments, where a similar value of qapp is obtained from the analysis of the ON and OFF gating current noise. We explain the different qapp for the ON and OFF test as deriving from the asymmetry of the energy profile of our model.

We imagined that the failure of the Markov model to describe the gating process of the Shaker channel, as simulated with our full Brownian model, could come from the following features of the channel. First, barriers are too low and highly asymmetric: the barriers between energy wells (measured in the forward direction) are all lower than 5kT (cf. Figure 4) and too highly asymmetric to be treated with the Markov model theory (Barcilon *et al.,* 1993)^14^. Second, gating charges on S4 are too close to each other: as we have learned from Figure 6, derived from the Simplified model, charges relatively close together tend to cross the pore simultaneously, thus enhance the gating current fluctuations and overestimate q_app_. We verified these propositions by looking at the modification of the variance-mean current plot produced by ad hoc modifications of these parameters of the model.

#### Dependence of the fluctuations on barrier height, friction coefficient and voltage

We first changed the height of the four barriers by adding or subtracting Gaussian-shaped energy components, to see whether the energy barriers present are not high enough for the full model to be consistent with Markov predictions (Figure 9Aa). As shown in Figure 9Ab, an increase in the energy barrier results in a lower q_app_ estimated from the variance-mean current plot (from 2.0, as shown in Figure 9B, to 1.75). These results tell us that the height of the barriers in the energy profile are not high enough to appropriately use Markov models. If the barrier heights are increased, the voltage sensor seems to activate more like a four step movement, each carrying one charge (i.e. q_app_ moves towards unity; Figure 9A). By contrast, reducing the energy barriers results in an apparent increase in the gating current noise and q_app_ (Figure 9Ac). We imagine that lower barriers increase the frequency of the gating charges passing simultaneously through the pore (see below), leading to higher current noise and q_app_.

**Figure 9.**
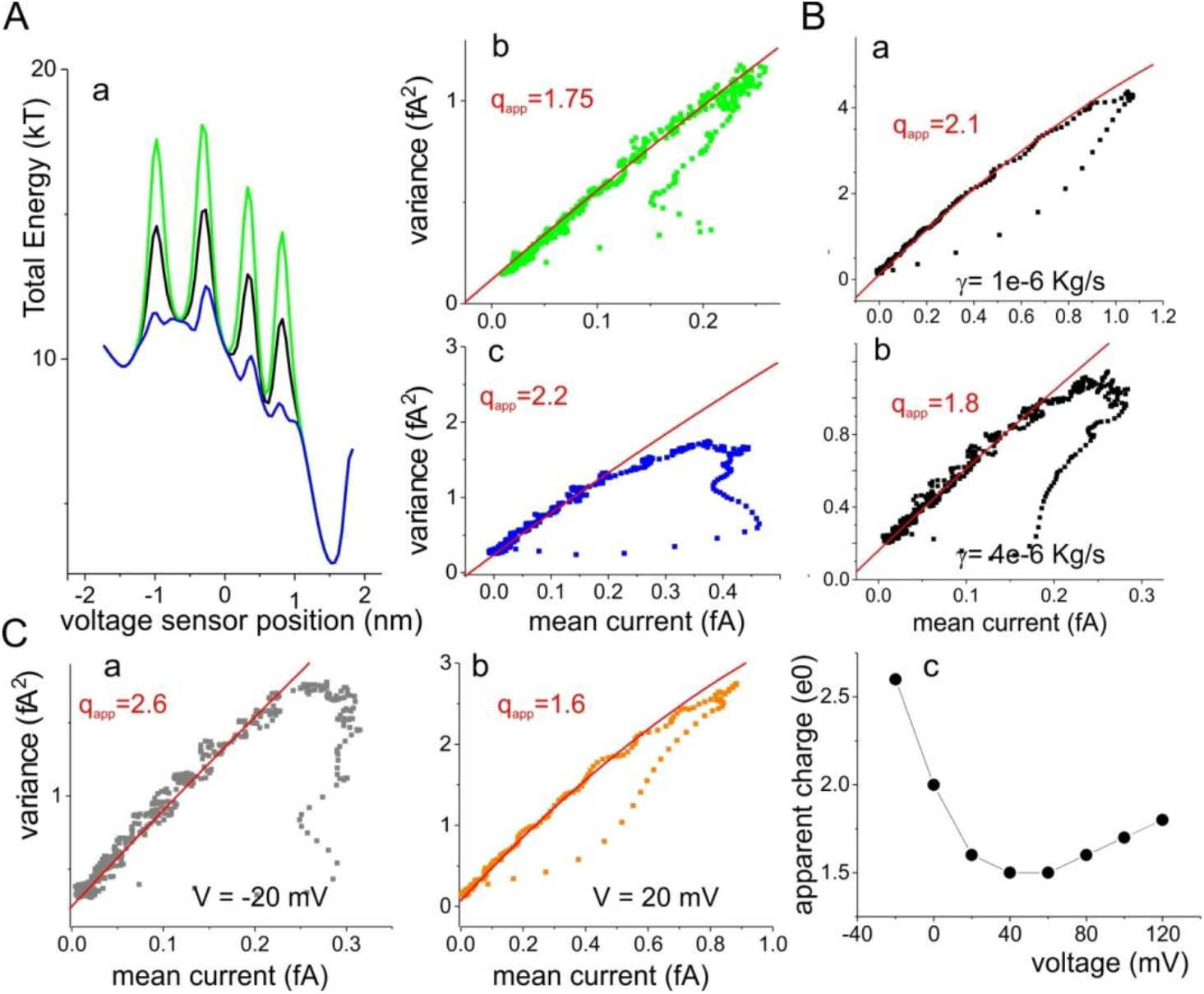
Dependence of the gating current fluctuations on the height of the energy barriers, the friction coefficient and the applied voltage. **A)** Panels b and c show the plots of variance vs mean current produced by altering the total energy profiles as shown in panel a (black line is the original energy profile of the Full model, green and blu lines are the altered energy profiles). An increase in the barrier height results in a decrease in the apparent charge. **B)** Panels a and b show variance-mean current plots obtained by altering the friction coefficient considered in the model, as indicated. **C)** Panels a and b show variance-mean current plots obtained by altering the applied potential, as indicated. Panel c plots the apparent charge obtained at various applied potentials.

With a voltage sensor moving in a non-Markovian regime (i.e., energy barriers not sufficiently high), we expect the gating current fluctuations to also depend on the frictional coefficient γ and on the applied potential V_m_, as shown with the simplified model in Figure 5 and Supplementary Figure 3. We indeed observed an inverse proportionality between q_app_ and γ (Figure 9B). As for V_m_ we observed a somewhat more complex situation, with a U-shaped q_app_ - V_m_ relationship (Figure 9C). Notably, an U-shaped dependence of the q_app_ has also been found experimentally (Rodríguez, Sigg and Bezanilla, 1998). As shown in Supplementary Figure 5 the significant increase in the q_app_ at moderately low voltages, that is not see in our Simplified model having only one gating charge (Figure 3), appears to come from the fact that the extreme position of the voltage sensor (the ones corresponding to the fully activated and fully deactivated position) are much more stable than the three intermediate position. In fact, if the voltage sensor is allowed to move through an energy profile where all states have approximately the same stability, the transient increase in the q_app_ at moderately low voltages disappears (Supplementary Figure 5).

Altogether, these results suggest that in a realistic model of voltage gating the voltage sensor dynamics produces a shot noise with features intermediate between a full Markovian (high barriers) and a drift-diffusion (no barrier) process.

#### Presence of multi-charge steps in the full model

We checked to see if the short inter-charge distance present in our model (and in real channels) produces a partial overlap of steps, leading to increased current fluctuations and q_app_. As shown in Figure 8A changes in the fluctuations were found changing the inter-charge distance of the four gating charges in both directions, with longer inter-charge distances giving an apparent charge closer to one.

Using the full model (with the original inter-charge distance), we also studied in detail the unitary events underlying the shot noise. We found a highly variable shape of the shots evoked by a depolarizing pulse to 0 mV, with the responses consisting of one to four well defined and distinct peaks, as shown in Figure 10B. From the analysis of 49 responses^15^, we found that those with three peaks were the most frequently observed (Figure 10Ca). In addition, we found that the amplitude and duration of the peaks appeared to decrease in the responses with a higher number of peaks (Figure 10C,b and c). From both the spatial trajectory of the voltage sensor and the time integral of the microscopic current shown in Figure 10B (central and bottom plots in each panel, respectively) it is possible to conclude that: i) in the responses with four peaks, each of them corresponds to the passage of one unitary charge through the gating pore, in other words, the passage of each charge is temporally separated from the passage of the other ones: activation process develops in four distinct steps (Figure 10Bb); ii) in the responses characterized by only one peak, the whole activation process develops in one single step that moves the four gating charges through the pore simultaneously, without an appreciable time lag between them (Figure 10Bc); iii) in the responses with two or three peaks, the integrals of current have values of 1, 2, or 3 e_0_, suggesting that 2 or 3 gating charges can simultaneously pass through the gating pore. In this case a single step in the activation process may actually represent the passage of 2 or 3 gating charges (Figure 10Ba,d). Figure 10D shows the charge amplitude histogram obtained from the 147 peaks detected in the 49 responses analyzed. The histogram displays four clearly defined peaks, approximately corresponding to 1, 2, 3 and 4 unitary charges, with frequencies of 0.68, 0.19, 0.09 and 0.04, respectively. This plot is a further evidence that the gating charges can cross the pore in rapid succession, in other words, in this case the voltage sensor moves by two positions skipping altogether a pause in the intermediate stable positions. Notably, a single shot analysis of the OFF gating current at −140 mV shows a lower number of multi-charge responses, especially those characterized by 3 and 4 unitary charges (data not shown). This may explain the reduced q_app_ obtained in these conditions (cf Figure 8F).

**Figure 10.**
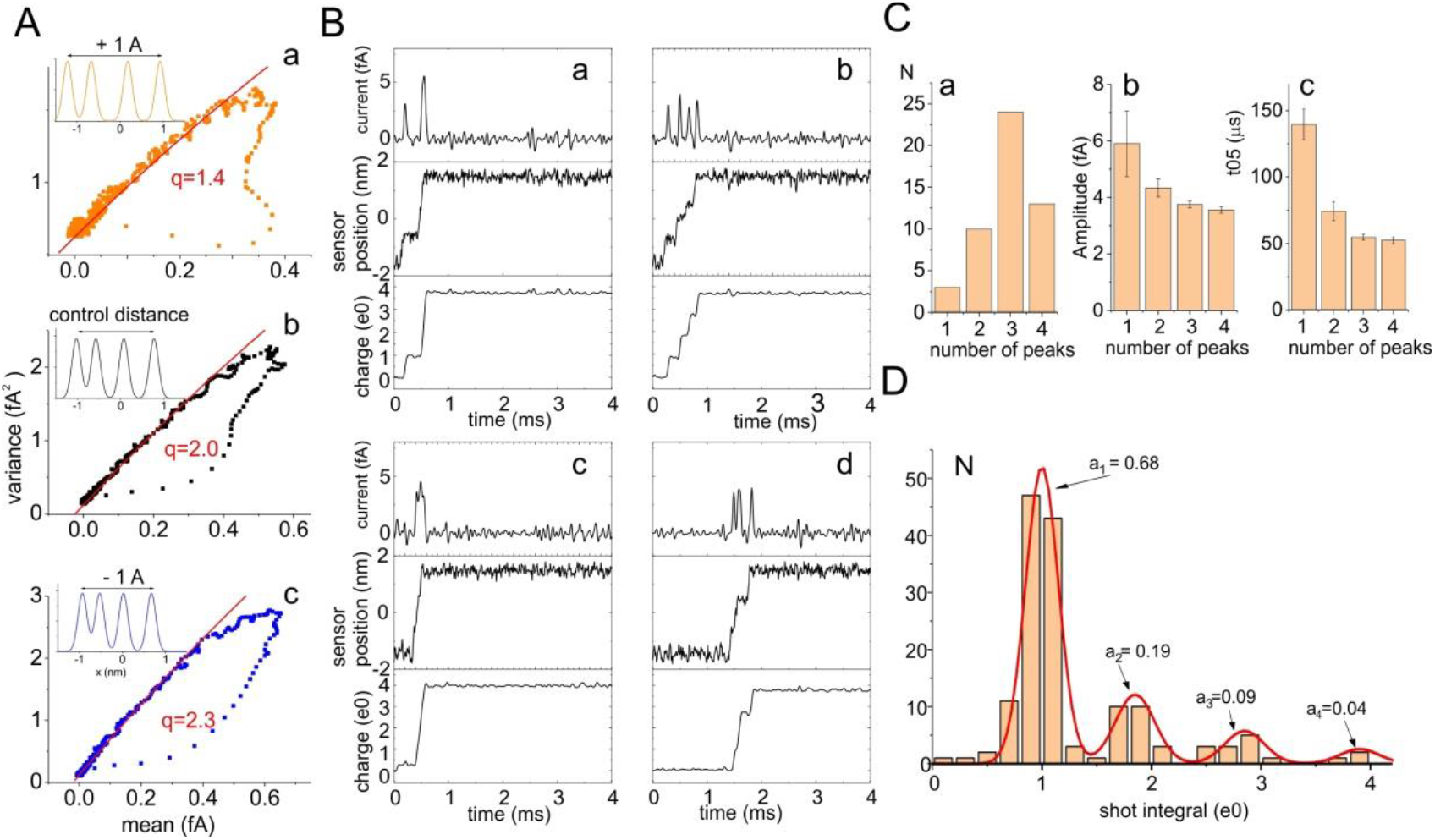
Evidence for multi-charge steps in the full model. **A)** Variance-mean current plots obtained with the Full model with the original setting (black data points) and with the inter-charge distance altered by −1 Å (blue) or +1 Å (orange). Insets show the corresponding gating charge distributions. **B)** Four typical microscopic current responses obtained at 0 mV are reported for the Full model, showing the presence of single (b) and multi-charge steps (a, c, and d). For each simulation the filtered current, the voltage sensor position, and the charge (time integral of the current) are shown. **C)** Panel a shows the number of responses with 1 to 4 peaks. Panels b and c show the mean amplitude and duration of the current peaks taken from responses with 1 to 4 peaks. **D)** Histogram of the charge carried by the current peaks, accumulated from 40 responses as those shown in panel B.

#### Dependence of the multi-charge steps on the applied voltage

As shown in Figure 9Cc, the q_app_ steadily increases with depolarization at voltages higher than 40 mV. In principle there are two possible reasons for this effect: i) as shown in Figure 5 and Supplementary Figure 3, in presence of relatively low energy barriers the shot current amplitude and duration are sensitive to the applied potential, and this may affect q_app_.; ii) a depolarization may increase the fraction of multi-charge steps, and this may reflect in a higher q_app_, as predicted by eqn. 14. We checked the validity of each mechanism by studying it in detail and comparing unitary responses produced by the Full model at +40 and +140 mV of applied potential. As shown in Figure 11A, at +40 mV responses were mostly characterized by 2 peaks, although responses with 1, 3, and 4 peaks were frequently observed. The corresponding amplitude histogram for the charge carried by the current peaks shows that at this applied potential the one-charge steps are the most frequent, while multi-charge steps are less frequent. By contrast at +140 mV the one peak response was domiant (Figure 11C). In this case events carrying four charges are much more heavily represented in the charge histogram (Figure 11D). These results are in accordance with a strong increase in the fraction of multi-charge steps with depolarization, that may well contribute to the increase in the q_app_ observed in Figure 9Cc and in the insets of Figure 11B and D).

**Figure 11.**
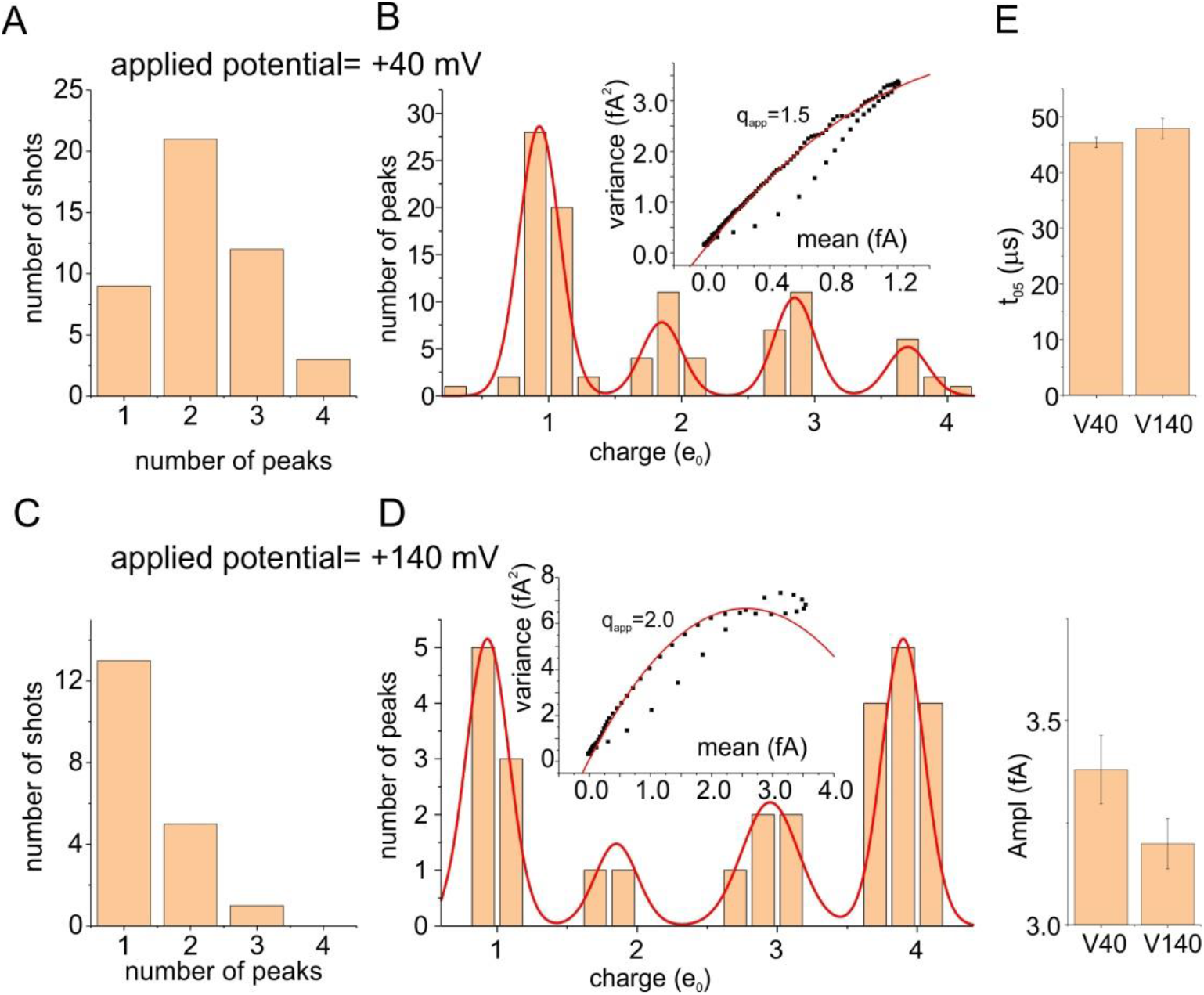
Voltage-dependence of the multi-charge steps in the Full model. **A) and C)** Number of responses with 1 to 4 peaks at applied voltages of +40 (A) and +140 (C) mV. **B) and D)** Histograms of the charge carried by the current peaks, accumulated from 40 responses, at applied voltages of +40 (B) and +140 (D) mV. Insets: Variance-mean current responses obtained at the same applied potential used in the corresponding histogram. **E)** Mean amplitude (Ampl) and dutation at half amplitude (t_0_._5_) of the current peaks carrying 1e_0_, at applied voltages of +40 and +140 mV.

In contrast there were no significant differences in the amplitude and duration of the current peaks carrying one charge (Figure 11E), indicating that the energy barriers encountered by the voltage sensor in the Full model do not allow the depolarization to significantly change the shape of the current peaks. In conclusion, of the two hypothesized mechanisms (change in shot current shape and change in the number of multi-charge steps) only the change in the fraction of multi-charge steps significantly contributes to the voltage-dependence of the estimated q_app_ at highly depolarized voltages.

Finally Supplementary Figure 6 shows an analysis of the frequency of all possible charge transitions, performed at four different voltages. It is clear that increasing the depolarization results in an increase in the probability of multi-charge transition. However it may be noticed that the first individual transition (S_0_->S_1_) appears also at relatively high voltages, in accordance with a relatively low charge initial step along the activation process previously noticed in (Sigg, Stefani and Bezanilla, 1994).

### Final conclusions

These results indicate that in our full model of voltage gating, and likely in the real Shaker channel, the energy barriers crossed by the gating charges are not high enough to allow the approximation of the dynamics of the voltage sensor by a Markov process. This implies that the gating current fluctuations and resulting q_app_ are not related in a simple and straightforward way to the amount of charge passing through the gating pore, as predicted by the Markov model. From the data shown here the current fluctuations and resulting q_app_ appear to depend mainly on the way gating charges cross the gating pore: as single charges, or as packages of varying multiple charges. These observations suggest that the interpretation of the variance-mean current plots and the resulting q_app_, obtained experimentally, needs to be made very cautiously^16^.

A simple four step Markov model appears unrealistic, as the charges interact with the medium and each other by electrical, steric, chemical and frictional forces and all those must be taking into account. In a few oversimplified but appropriate words ‘everything interacts with everything else’.

## Supplementary Data

**Supplementary Figure 1.**
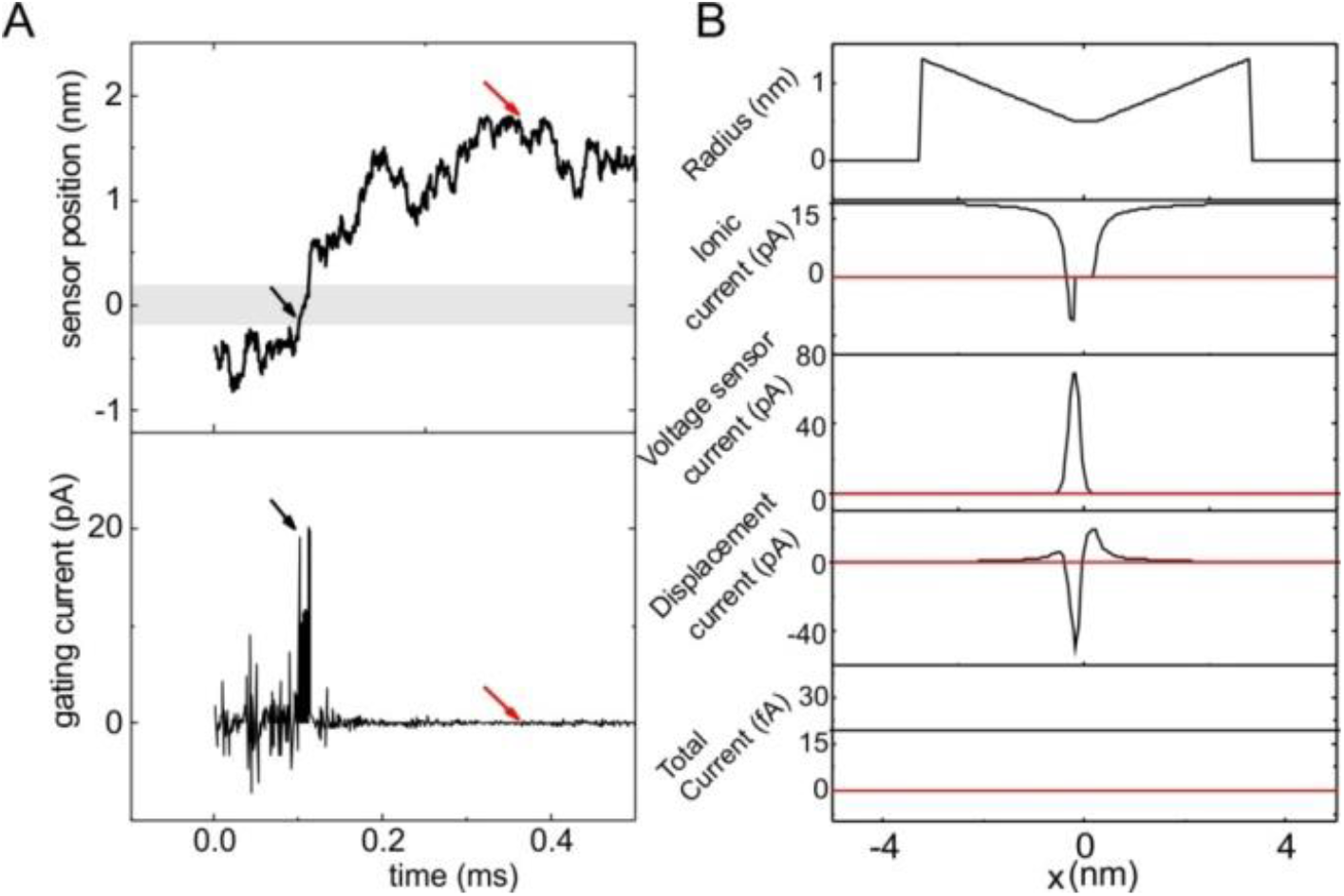
Test of the total current conservation in the Simplified model. **A)** Time dependent position *(top)* and unfiltered gating current *(bottom)* obtained in a simulation with the Simplified model. Friction coefficient and energy profile are the same used in Figure 4A of the main paper. Black and red arrows indicate the points in time chosen to verify the total current conservation. **B)** Spatial profile of (from top to bottom): i) radius of the voltage sensor domain; ii) the ionic current, assessed as 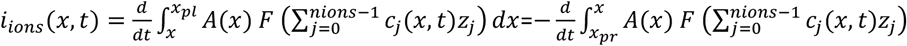 where *x_pl_* and *x_pr_* are the left and right extremes of the gating pore, *F* is the Faraday constant, and *Z_j_* is the valence of ion j. iii) the current carried by the voltage sensor, assessed as 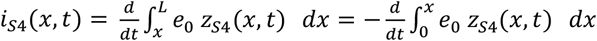, where *Z_S4_(x, t)* is the charge density profile of the S_4_ segment; iv) the displacement current, defined as 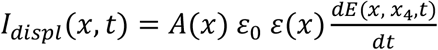, where *E(x, x_4_, t)* is the electric field, for which we have explicitly indicated the dependence on the spatial dimension, time, and position of the voltage sensor x_S4_, *A(x)* is the area at position x, ε_0_ is the permittivity of free space and *ε(x)* is the position-dependent dielectric constant; v) total current, defined as *i_tot_(x, t)* = *i_ions_(x, t)* + *i_S4_(x, t)* + *i_displ_(x, t)*. Red and black lines refer to the profiles assessed at the two time points indicated with the arrows of same colors in panel A. For a more extensive treatment and derivation of the current conservation and of the mathematical forms of the various current see Supplementary material in Catacuzzeno et al., (2019).

**Supplementary Figure 2.**
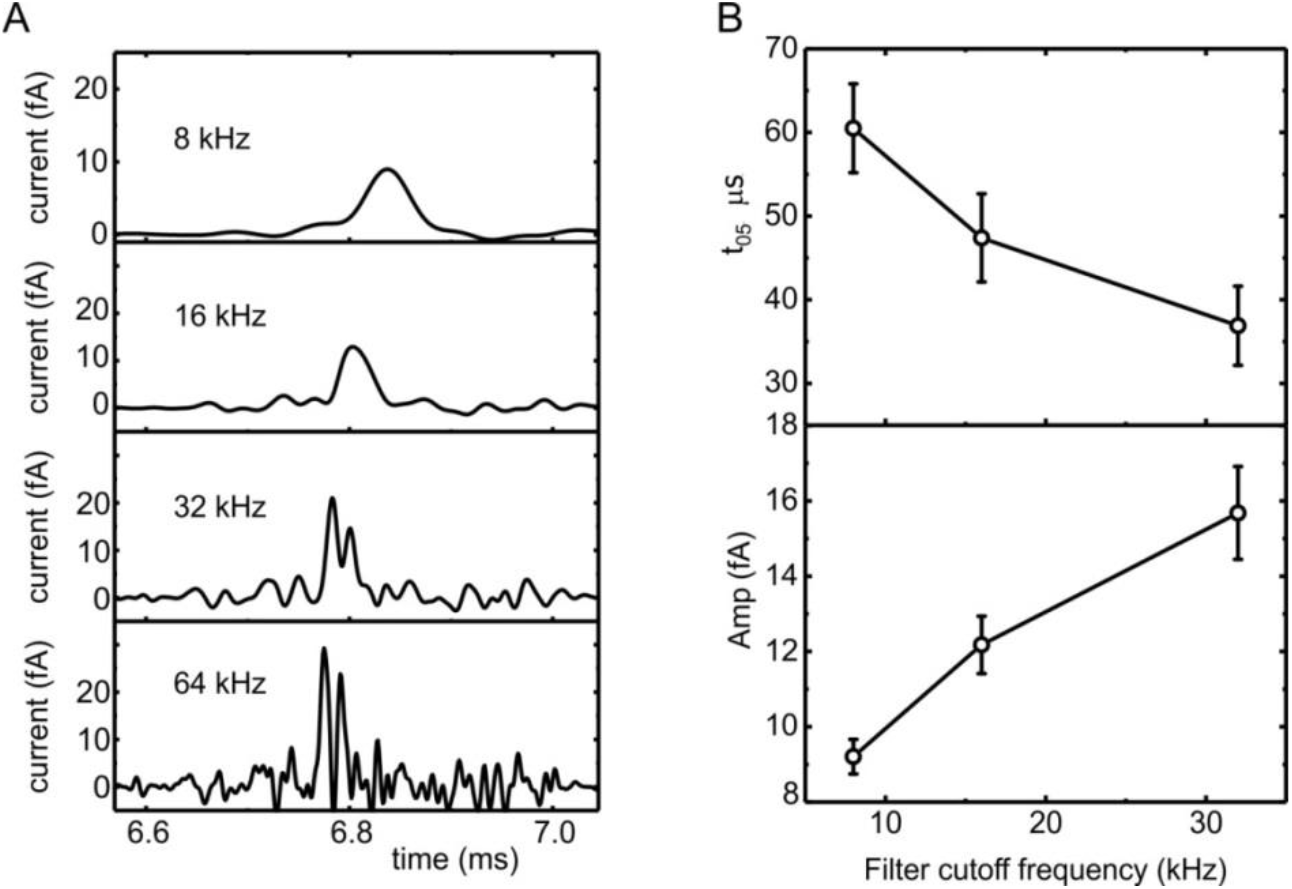
Effects of filter cut-off frequency on the shape of the current shot. **A)** Current shot simulated with the simplified model, no energy barrier (same parameters of Figure 2C of the main paper), filtered with an 8-pole Bessel filter at varying cut-off frequency (indicated). **B)** Plot of the amplitude (Amp) and duration of the current shots at half amplitude (t_0_._5_), estimated at different filter frequencies.

**Supplementary Figure 3.**
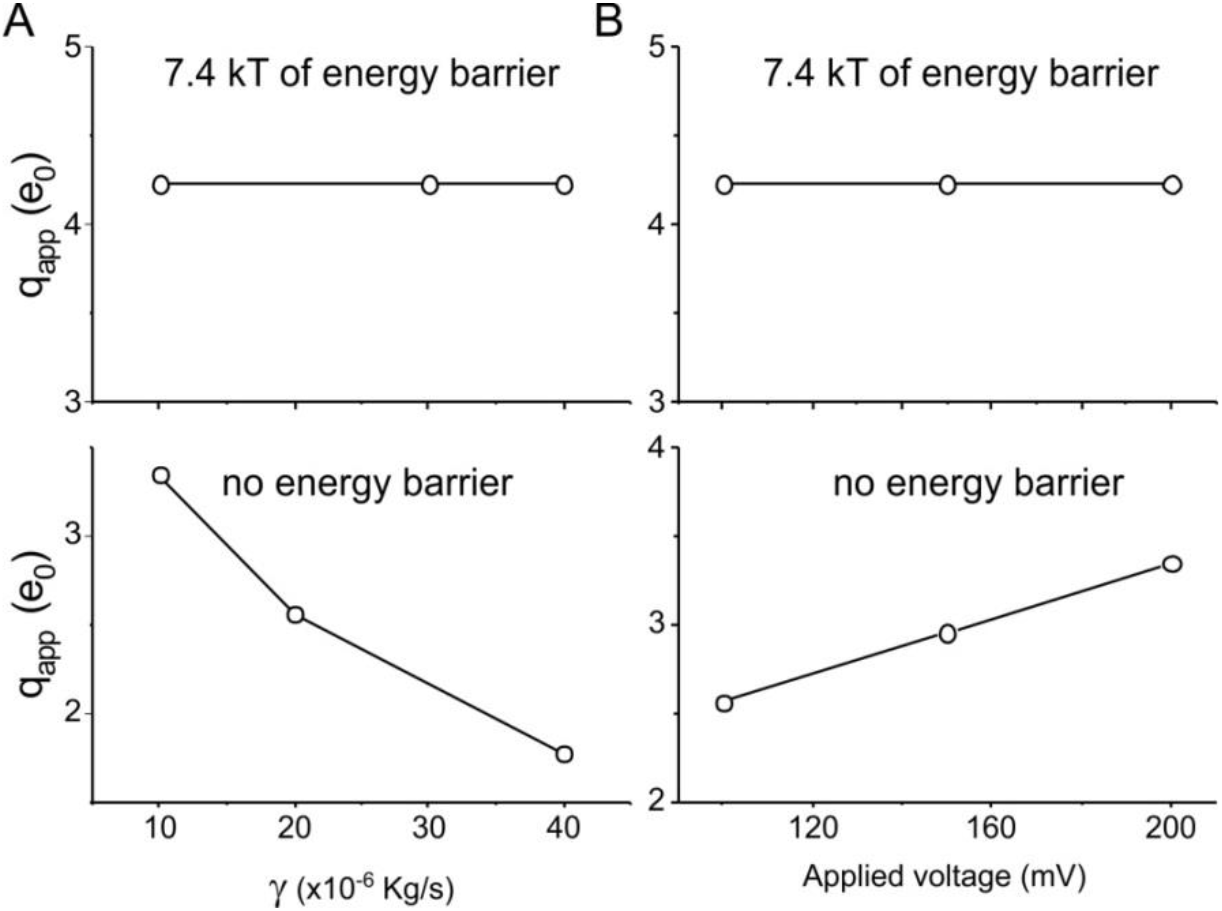
Differential dependence of the apparent charge on the friction coefficient and applied voltage. Plots of the apparent charge estimated from the Simplified model with (7.4kT, upper plots) and without (lower plots) an energy barrier, at varying friction coefficient (A) and applied potentials (B). The plots clearly show that the apparent charge depends on the two parameters in the no-barrier case, but not in the high barrier case.

**Supplementary Figure 4.**
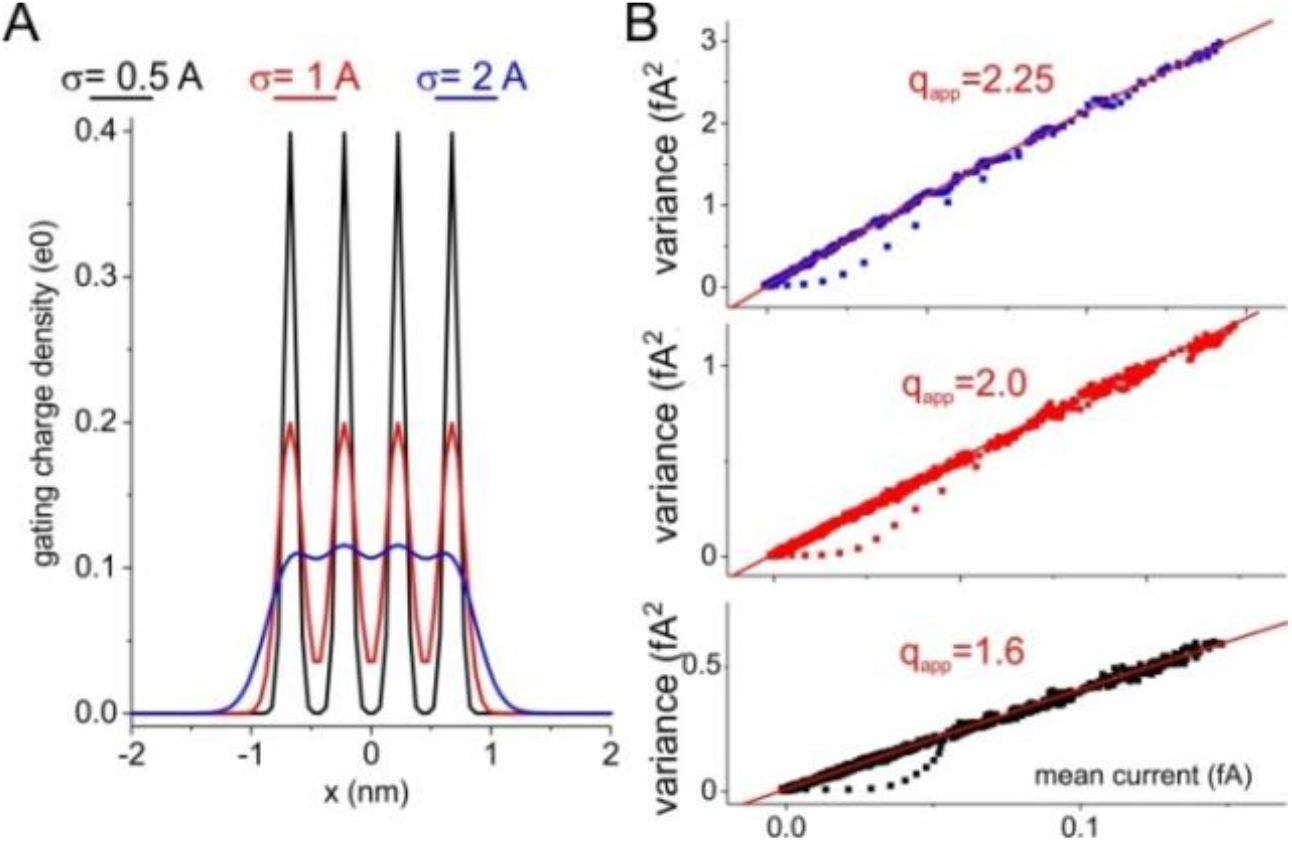
Dependence of the apparent charge on the spreading of the gating charges. Panel B shows variance-mean current plots obtained from the Simplified model, including four gating charges at an inter-charge distance of 5 Å. In the three simulations the standard deviation of the normal distribution used to distribute the gating charges was varied, as indicated in panel A.

**Supplementary Figure 5.**
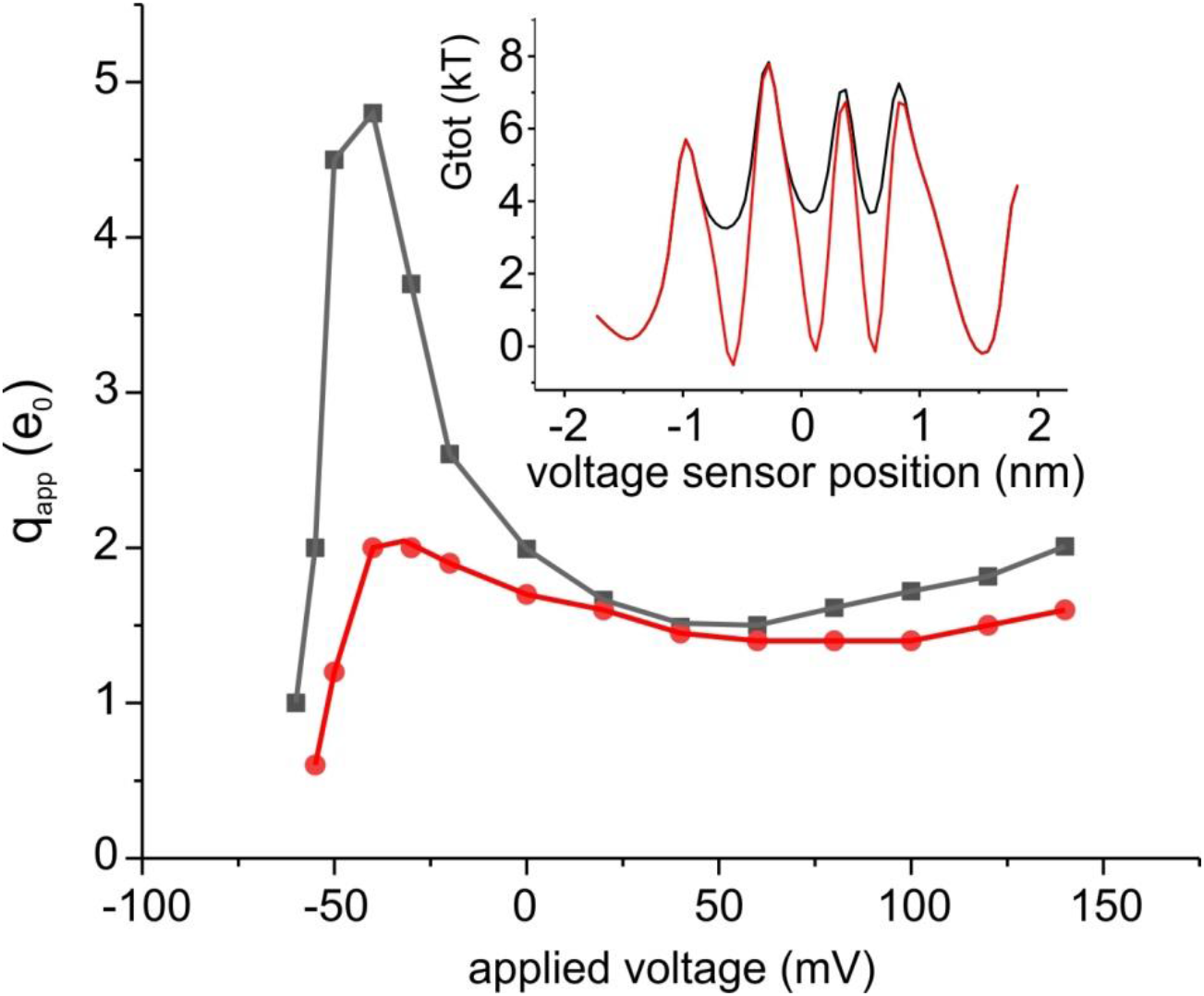
Plot of the apparent charge obtained at various applied potentials for the Full model (black symbols) and for a model where the energy profile encountered by the voltage sensor was appropriately modified to give about the same stability to all states of the voltage sensor (red symbols). The inset shows the energy profile of the Full model at −40 mV (black profile), and the modified energy profile (red) at the same applied voltage.

**Supplementary Figure 6.**
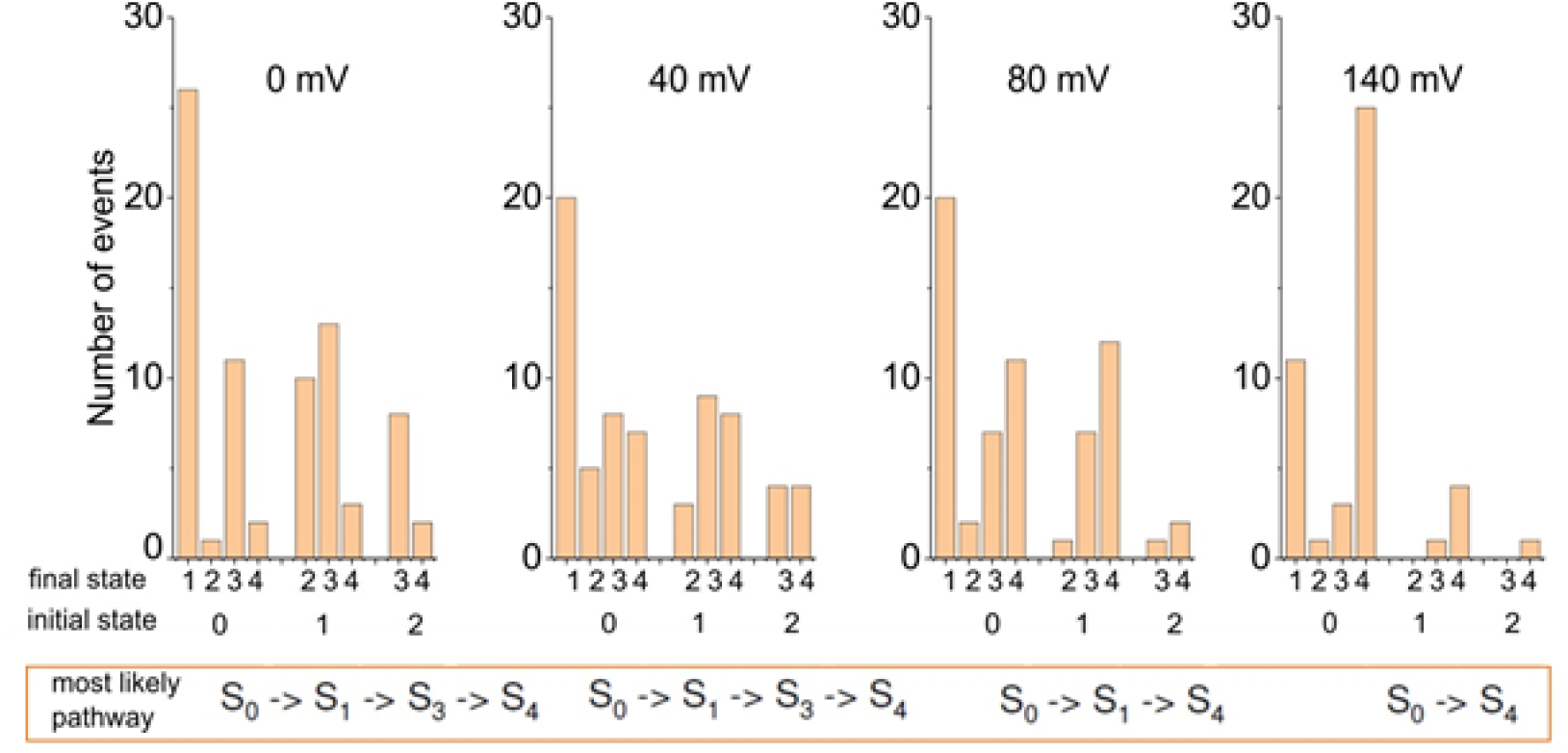
The plots were obtained from the analysis of 40 responses obtained with the Full model, at four different applied potentials. Each bar represents the number of times we observed the passage of the voltage sensor (as evaluated from the gating charge change) from the indicated initial state to the indicated final state. The state numbering corresponds to the number of gating charges found in the right vestibule. At the bottom, framed with an orange line, are shown the most represented pathways followed by the voltage sensor, starting from state 0 and following the most probable event. These data suggest that i) the state 2 has a very low probability of being occupied; ii) the step 0 -> 1 is the most represented, even at relatively high voltages (except for at +140 mV).

### Interpretation of the variance to mean current plot for a signal with multiple types of shot

In this derivation, we followed the formalism of Conti and Stuhmer (1989), by considering the case of the simultaneous presence of multiple types of shots, each carrying a different amoint of charge, contributing to the signal. Consider a signal composed of N types of shot events carrying different charges q_j_ and represented at a frequency a_j_, where j=1.,.N. Let p(t_j_) be the probability density that the shot carrying a charge q_j_ begins at time t_j_. If the signal is filtered by a Gaussian filter with a cutoff frequency of B, having a Gaussian response function 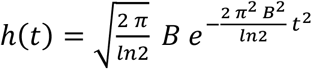 the average current observable in repeated trials will be:

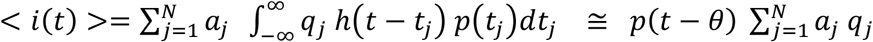

Where the last approximation is valid if the signal varies slowly as compared to h(t). The corresponding mean squared current observed in repeated trials will be:

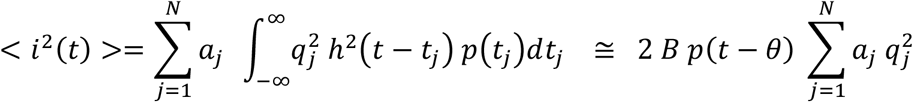

where we have considered that 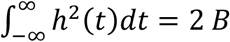 for a Gaussian filter, and as a first approximation also for an 8-pole Bessel filter.

The variance of the signal will thus be

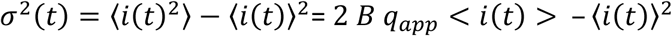

where 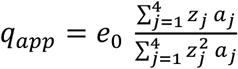

### Numerical solution of the PNP system

#### Discretization of the flux conservative equation

Ions in the baths and vestibules were subjected to electro-diffusion governed by the following flux conservative equation:

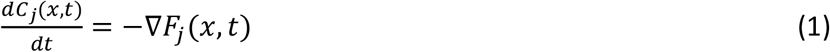

that for our mono-dimensional model may be discretized following Figure S6, obtaining the following eqn.

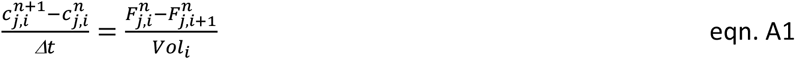

where 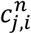 represent the concentration of ion j in the i^th^ volume element at time n Δt, Δt is the time-step, 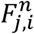 is the flux (moles per unit time) of ion j from volume element i-1 to volume element i, and Vol_i_ is the volume of volume element i. The ion flux follows the Nernst-Plank equation:

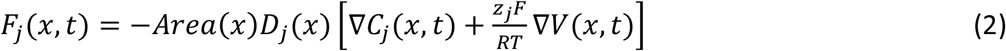

That in our case may be discretized in the following way:

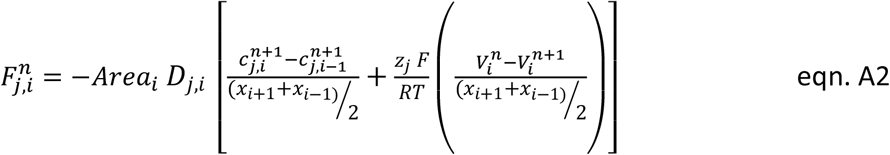

Where *Area_i_* is the surface of the left border of the i^th^ volume element, *D_j,i_* is the diffusion coefficient of ion j inside the i^th^ volume element, *x_i_* is the position of the left border of volume element i^th^, z_j_ is the valence of ion j, F is the Faraday constant, R is the universal gas constant, T is the absolute temperature and 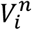 is the electric potential profile in the i^th^ volume element at time n Δt.

**Supplementary Figure 7.**
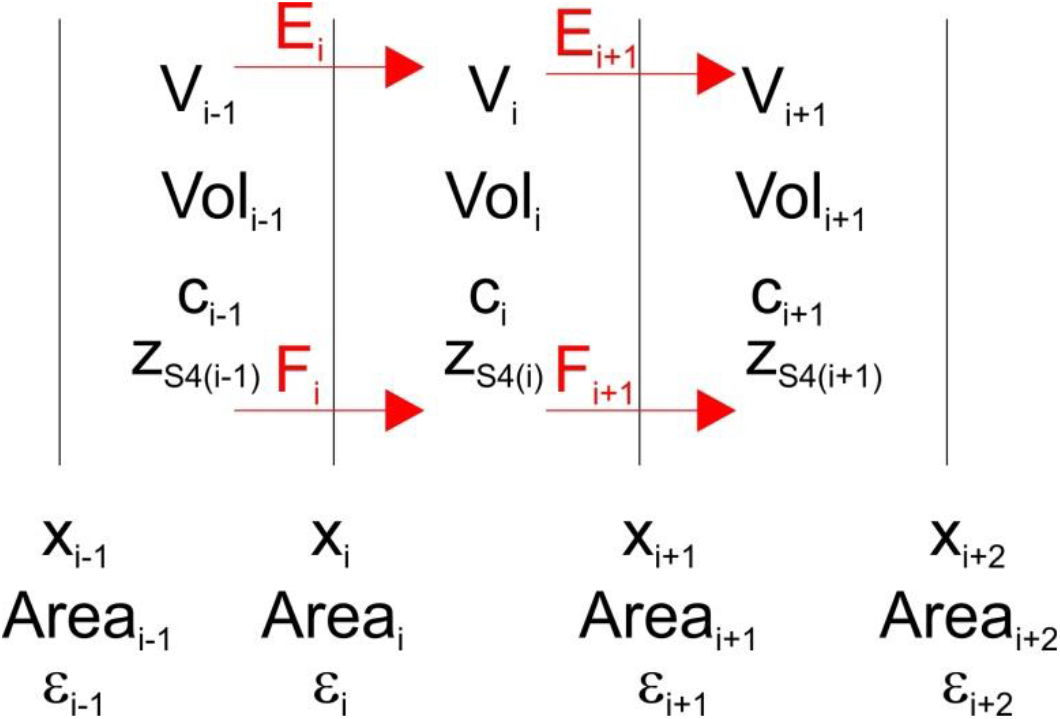
Discretization of the spatial domain in our model

Equations A1 and A2 may be combined to give the following equation

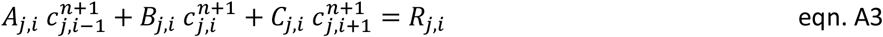

Where

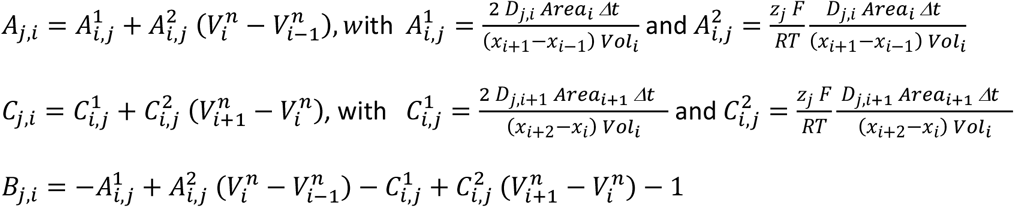

Equations A3 form a linear system of N-2 equations, with i=1,........, N-1. They can be coupled with the boundary conditions imposing a constant ion concentration at the left and right boundaries

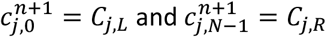

to obtain a set of N linear equation for the N unknown 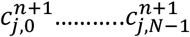 that may be solved with an algorithm for tridiagonal systems as given in Press et al., 1992, thus recovering the ion concentration profiles at each timestep *Δt*.

#### Discretization of the Gauss law (Poisson equation)

Gauss law of electrostatics (that represent the integrated form of the Poisson’s equation) relates the flux of the electric field out of a closed surface to the net charge existing inside

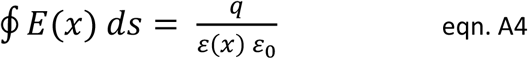

Where ∮ is the integration over a closed surface, E(x) is the electric field, q is the charge contained inside the closed surface, *ε_0_* is the permittivity of free space and *ε(x)* is the relative dielectric constant. Applying gauss law to the i^th^ volume element we obtain:

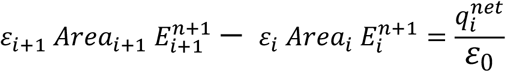

Where ε_i_ is the relative dielectric constant across the left boundary of the volume element, and 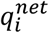 is the net charge inside the i^th^ volume element, given by

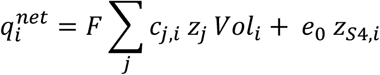

Where *e_0_* is the elementary charge and *z_S4,i_* is the amount of gating charge inside the i^th^ volume element. Considering that

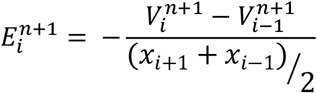

We obtain the following linear system of equations:

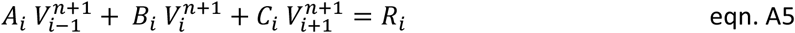

Where

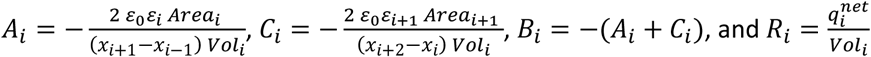

Equations A5 form a linear system of N-2 equations, with i=1,........, N-1. They can be coupled with the boundary conditions imposing a known applied potential at the left and right boundaries

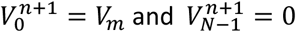

to obtain a set of N linear equation for the N unknown 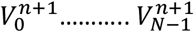 that may be solved with an algorithm for tridiagonal systems as given in Press et al., 199,2 thus recovering the electrostatic potential profile at each timestep *Δt*.

#### Steady state solution of the PNP system

In our model we assume that ions relaxes much faster that the voltage sensor. This means that for each given position of the voltage sensor the ion concentration and electrostatic potential profiles assume instantaneously their equilibrium values. In our computations these equilibrium profiles were found by iteratively solving eqn A3 and A5 using a timestep of 0.2 ns, until finding equilibrium, that was defined as the situation in which the maximum relative change in the electrostatic potential within the all spatial profile obtained in the iteration was lower than 10^−8^.

## Footnotes

1 Simulations show that those restrictions are more limiting than often thought: a large barrier for example must be both large and symmetrical to ensure the approximation. The asymmetry between starting state and ending state must be small compared to the barrier height (Barcilon *et al.*, 1993).

1 Although a travelling distance for the voltage sensor of 36 Å may seem too large, several points need to be considered. First in our Full model this travelling distance allowed the movement of the voltage sensor by the distance separating R1 from K5 (28 Å) assessed by taking the charge-to-charge distance in the 3D structure of the VSD (Catacuzzeno and Franciolini, 2019b). Considering that this distance corresponds to the passage through the pore of the first four gating charges (R1-R4), those relevant to gating, the average distance each of them has to travel along the longitudinal axis to cross the gating pore is ca. 7Å, an amount in accordance with the available literature. Indeed, although the current consensus (Vargas, Bezanilla and Roux, 2011) indicated a maximal vertical motion of 10 Å, the simulations showed that the sensor moved along a path that was about 40° off the membrane normal, thus giving a motion along the main axis of the voltage sensor of 12.5 Å. We also need to consider the position of the voltage sensor at rest. It is now clear that there are more than one ‘resting’ state, whose specific occupancies depend on the level of hyperpolarization, which is different in different studies. To this respect, the model of (Vargas, Bezanilla and Roux, 2011) took as ‘closed state’ the one that is most populated at moderate negative voltage (the ‘penultimate resting state’, as it was called by (Lin *et al.*, 2011)), as opposed to the resting state reported, for instance, by (Tao *et al.*, 2010) and (Delemotte *et al.*, 2011), with R1 in the catalytic center, which is reached only at high negative voltages, that we can call the “deep resting state”. This state has been also proposed by (Henrion *et al.*, 2012) for Shaker channels, using a combination of modeling and experiment work, and named C4, in which R1 lies below F290 and interacts with E293. The model proposed suggests an S_4_ movement of at least 12 Å to pass from the active state, O, to the C3 state, while the deeper resting state C4 of the VSD could be reached with an S4 movement of ∼17 Å (Henrion *et al.*, 2012). Now, even taking 17 Å as a reliable guess of the distance travelled by the voltage sensor for full VSD activation, our data are still significantly higher than this. The gap in our view derives in large part from the fact that in our model the charge is glued on the S_4_ segment, instead of extending out by 6-7Å (the arginine length), which decreases by about another 4-5 Å, considering that the first and last gating charges – R1 and R4 – are thought to be tilted by 30-45° from S_4_ normal, and in opposite way, when bound to the catalytic center, containing in this way the voltage sensor movement. All this considered, the difference between our data and literature can be considered minor.

3 For simplicity, from here on baths and vestibules on either side of the gating pore will be considered as a single environment and referred to as ‘bath’

4 This is a verbal statement of the continuity equation in one dimensional unbranched systems which is equivalent to saying that total current is the same everywhere in such a circuit. The mathematical statement is important because it is precise as no verbal statement can be (Eisenberg, Kłosek and Schuss, 1995).

5 Notice that the chemical component of the energy might also include the electrostatic interaction between the gating charges as counter-charges present on the vestibules. A potential profile can be separated into components in many ways. Each separation takes on physical meaning when its boundary conditions and functional dependence (e.g., defining differential equation) is specified. It is very important to verify that the sum of the components, computed with its boundary conditions, actually add up to the original function. This is not at all obvious because the boundary conditions may differ. The classical superposition theorem is sometimes used without considering the role of boundary conditions that are a mathematical necessity. Practically speaking the boundary conditions also change the numerical value of many results by a great deal.

6 We call this charge apparent since, as we will see, it does not always assume the value corresponding to the gating charge moving in more physically realistic models.

7 We verified that the small overestimate of the gating charge originates from the ringing property of the Bessel filter we used, and could be mostly eliminated using a Gaussian filter.

8 Varying the barrier height also changes the time course of the gating currents. For this reason the friction coefficient γ was changed in the various simulations, to keep the time course of gating currents approximately the same.

9 Actually, because in our model the charge is not a point charge (and this is a realistic situation, due to the atomic dimension of the gating pore) the shots produced are not expected to be exactly square current pulses, but to have a not-instantaneous rising and decay time course that for simplicity we have not considered in the analysis presented.

10 Indeed, as previously pointed out (Sigg et al., 1994), in a multi-step Markovian process the q_app_ resulting from the variance-mean current plot cannot be larger than the largest charge carried by a single step. A multi component barrier model that includes interactions (i.e., correlations), as required by the Maxwell equations in the ion movements might have quite different behavior (Eisenberg, 2020).

11 In this study we used relatively high energy barriers, of ~4.6 kT, see Figure 9C.

12 i.e., two or more consecutive charges pass through the pore in rapid succession without the voltage sensor dwelling in the energy well in between for a visually perceptible or measurable amount of time, and in this case it is viewed as one step

13 The energy profile in the full model is not assumed as in the simplified model, but self-consistently assessed using the Poisson equation. We add that a spring component is also present in this model to reproduce the charge vs potential curve of experiments (Horng *et al.,* 2019; Catacuzzeno, Sforna, Franciolini and R. S. Eisenberg, 2020).

14 While Markov models have evident appeal, it must be remembered that they have no logical significance of their own: the field equations of physics (Morse, P. M. C., 1953; Arfken, G. B., H. J. Weber, 2013; Fl, Landau and Ml Lifshitz, 2013) cannot be cast as Markov processes, because they involve space and time dependence beyond that possible in the Markov structure. An important case is discussed in (Eisenberg, 2020).

15 These were selected from a computation of 60 responses, by discarding trials where the current peaks were not very well defined because of the superimposition of negative peaks coming from short backward movements of the voltage sensor.

16 Our data somehow fulfill the idea originally put forward by (Crouzy and Sigworth, 1993) that, because “the filter bandwidth is limited, a rapid succession of small charge movements can become indistinguishable from a single large charge movement”. As a consequence, “the figure of 2.3eo obtained for the two-state model might not reflect the true size of the elementary charge movements, because it could represent the sum of charge movements that occur relatively rapidly in the gating process of the channel”.

